# Discovery of a Plant Pictet-Spenglerase with R-Stereoselectivity

**DOI:** 10.64898/2026.02.05.704076

**Authors:** Clara Morweiser, Sarah Heinicke, Sarah E. O’Connor, Maite Colinas

## Abstract

The class-defining monoterpenoid indole alkaloid (MIA) scaffold strictosidine is generated by condensation of tryptamine and secologanin by the Pictet-Spenglerase strictosidine synthase (STR). All previously characterized STR orthologs are strictly 3*S*-stereoselective. Here, we report that the medicinal plant species *Pogonopus speciosus* (Rubiaceae) accumulates the 3*R* epimer vincosidic acid produced by a an ortholog of STR. This ortholog, named here Epi-STR, exclusively produces the 3*R* configuration and is capable of accepting both secologanic acid and the methylester secologanin as aldehyde substrates. Using comparative phylogenetic and structural analyses, we determine the amino acid residues that confer stereoselectivity. Through rational design of reciprocal amino acid substitutions, we achieved switches in stereoselectivity in a canonical STR and in Epi-STR. The stereoselectivity of the engineered mutants is also dependent on the identity of the aldehyde substrate. Notably, previous and extensive engineering efforts have failed to switch the stereo-selectivity of STR. Therefore, this discovery now allows cost-effective epimer-pure access to the *R*-epimer, and offers mechanistic insights into the enzyme stereoselectivity of this important reaction. This work also highlights the importance of phytochemical analyses of poorly described plant species.

---

The Pictet-Spengler reaction, which is the condensation between an arylethylamine and an aldehyde or a ketone followed by ring closure to yield a tetrahydro-β-carboline or a tetrahydroisoquinoline moiety, is a reaction frequently used in chemical synthesis ^[1]^. In plants, the central scaffolds of diverse alkaloids are formed by a Pictet-Spengler reaction including monoterpenoid indole alkaloids (MIAs), ipecac alkaloids, benzylisoquinoline alkaloids (BIAs), and phenylethylisoquinoline alkaloids (PIAs) (reviewed by ^[2]^). While non-enzymatic Pictet-Spengler reactions may occur in certain plants, these reactions are most commonly catalyzed by Pictet-Spenglerase enzymes ^[3]^. Only two plant enzyme families are known to have evolved Pictet-Spenglerase activity: the pathogenesis-related 10/Bet v1 (PR10) protein, which catalyzes the Pictet-Spengler reaction between dopamine and an aldehyde to yield a tetrahydroisoquinoline product (BIA and PIA biosynthesis), and strictosidine synthase (STR), which uses tryptamine and an aldehyde to yield a tetrahydro-β-carboline (MIA biosynthesis) product ^[3d, 4]^. Many MIAs are medically important due to their anticancer (vinblastine, camptothecine), antimalarial (quinine), antihypertensive (ajmalicine, reserpine), or neuroactive (mitragynine, ibogaine) activities ^[5]^. The class-defining MIA scaffold is formed by the Pictet-Spengler reaction of tryptamine **1** with the secoiridoid secologanic acid **2** or the methylester secologanin **3** (Figure 1a). While this reaction can occur spontaneously and non-stereoselectively in solution under strong acidic conditions, the STR enzyme is strictly stereoselective and exclusively produces the 3*S* configuration strictosidinic acid **4S** or strictosidine **5S**, respectively (Figure 1a). The natural secoiridoid substrate in the Cornales order (including *Camptotheca acuminata*) is secologanic acid **2** while plants in the Gentianales order (including *Catharanthus roseus*) use the methylester secologanin **3** ^[6]^. Consistent with the strict *S*-stereoselectivity of STR, the alternative 3*R* Pictet-Spengler products vincosidic acid **4R** or vincoside **5R** do not usually accumulate in plants. Changes in stereoselectivity of STR have only been observed for *Ophiorrhiza pumila* STR (OpSTR) when non-native, small aldehydes are substituted for the secoiridoid aldehyde substrate ^[7]^. In some MIAs the C3 stereocenter is switched from *S* to *R* downstream of strictosidine **5S** by the activity of epimerization enzymes ^[8]^. STRs have long been a target for enzyme engineering with the aim of broadening the substrate specificity and/or altering stereo-selectivity. However, despite extensive engineering efforts, production of **4R** or **5R** has never been achieved ^[4c, 7, 9]^. Here, we report that the medicinal *Rubiaceae* plant *Pogonopus speciosus* harbors an *R*-selective STR ortholog, here named Epi-STR, which is responsible for the exclusive production of the 3*R* epimer (vincosidic acid **4R**) found in this plant. We further identify specific amino acid residues that confer stereoselectivity, providing a mechanistic basis for this inverted stereochemical preference.

**Figure 1.**
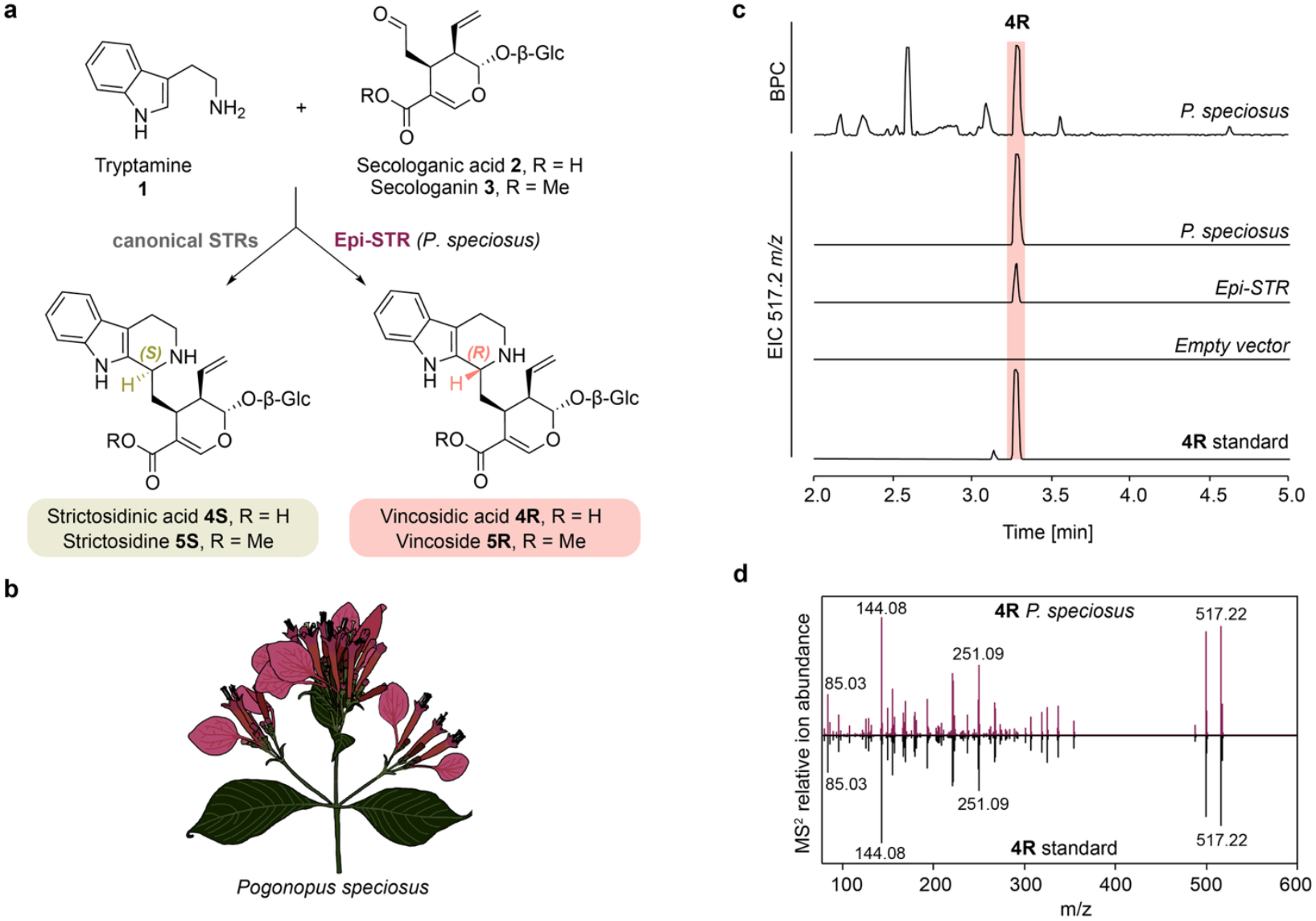
*P. speciosus* produces vincosidic acid **4R** catalyzed by the *R*-selective Pictet-Spenglerase Epi-STR. **a**, Scheme showing reactions of canonical STRs and *P. speciosus* Epi-STR reported here. Tryptamine **1** is coupled with a secoiridoid, either secologanic acid **2** or the methylester secologanin **3** to the *S*-epimer by canonical STRs or the *R*-epimer by Epi-STR. Note, that the common native substrate of canonical Gentianales STRs is secologanin **3** whereas it is secologanic acid **2** for Epi-STR. **b**, Representative drawing of a *P. speciosus* inflorescence. **c**, Liquid chromatography-mass spectrometry (LC-MS) traces from extracts from *Pognopus specious* young leaves, *Nicotinana benthamiana* leaf disks expressing *Epi-STR* or empty vector control and fed with tryptamine **1** and secologanic acid **2**, and authentic standard of vincosidic acid **4R** (see methods). BPC, base peak chromatogram. **d**, MS^2^ fragmentation pattern of *P. speciosus* and authentic standard peaks confirming the identity of the peak as vincosidic acid **4R**.

*P. speciosus* is known to produce ipecac alkaloids with antimalarial activity ^[10]^. Although MIAs have never been reported from this species, when we analyzed the alkaloid content of young leaves through liquid chromatography-mass spectrometry (LC-MS) we detected a high accumulation of vincosidic acid **4R**, as verified by comparison to an authentic standard (Figure 1b-d). We performed IsoSeq sequencing on a pooled RNA sample of a mixture of tissues and constructed a de novo transcriptome assembly (Figure S1). Blasting of the *C. roseus* STR (CrSTR) amino acid sequence against the assembled transcriptome resulted in the identification of a single ortholog with 51% sequence identity to CrSTR ^[11]^. Transient overexpression of this ortholog in *Nicotiana benthamiana* leaves followed by feeding of the substrates tryptamine **1** and secologanic acid **2** to cut leaf disks led to the exclusive formation of vincosidic acid **4R** (Figure 1c; S2). Although vincoside **5R** could not be detected in *P. speciosus*, Epi-STR also catalyzed formation of **5R** when secologanin **3** was provided as a substrate (Figure 3i). In contrast to other related plant species, we did not identify an ortholog of the methyl transferase that converts this acid moiety to the methyl ester (loganic acid methyltransferase (LAMT)), which explains why only secologanic acid **2** is available as a substrate in *P. speciosus* ^[12]^. To characterize the catalytic activity of Epi-STR, we carried out steady-state kinetics (Figure S3). We determined a K_m_ value of 636.8 µM for tryptamine (k_cat_ of 0.1 s^-1^), which signifies a significantly lower catalytic activity of Epi-STR compared to the canonical CrSTR from the Apocynaceae plants *C. roseus* (4 µM; 5.1 s^-1^) and *Rauvolfia serpentina* RsSTR (6.2 µM; 1.3 s^-1^) ^[4c, 13]^. K_m_ values for secologanic acid **2** and secologanin **3** are estimated at > 5 mM but were too high to be reliably measured, and are significantly higher than the K_m_ values reported for secologanin **3** for CrSTR (40 µM), RsSTR (39 µM) and OpSTR (21 µM) ^[4c, 9e, 13]^. Although Epi-STR appears to have comparably low catalytic activity, vincosidic acid **4R** is among the highest accumulating metabolites in *P. speciosus* young leaves (Figure 1c). Of note, secologanin **3** concentrations exceeding 100 mM were previously measured in *C. roseus* leaf single cells ^[14]^. It is therefore plausible that similar cellular concentrations of secologanic acid **2** may occur in *P. speciosus*, explaining the efficient production of vincosidic acid **4R** despite the high K_m_ value of this enzyme.

Next, we compared the active site of Epi-STR with those of canonical STRs through a combination of structural and phylogenetic analyses (Figure 2). STRs from the Gentianales form a sister clade to the structurally related strictosidine-synthase-like enzymes (SSLs), which ubiquitously occur in all plants, but whose functions are unknown (Figure 2a, S4) ^[15]^. Rubiaceae members frequently contain two STR paralogs, but in *P. speciosus* Epi-STR was the only observed ortholog (Figure 2a). An overlay of an Epi-STR Alphafold3 model with reported crystal structures of the Apocynaceae RsSTR (PDB code: 2v91) and the Rubiaceae OpSTR (PDB code: 6s5j) shows that the six-bladed β-propeller fold is entirely conserved (Figure 2b) ^[7, 16]^. Previous structure-activity studies on canonical STRs, in particular RsSTR and CrSTR, identified active site residues (Figure 2c) ^[4c, 9e, 13, 16]^. A sequence alignment of canonical STRs and Epi-STR revealed that nine active site residues at the pocket entrance that appear to interact with the secoiridoid moiety are substituted in Epi-STR. In contrast, six residues that likely interact with the tryptamine **1** substrate are conserved, including the glutamate residue that is essential for catalysis (Figure 2a,c) ^[4c]^. Mutating this glutamate in Epi-STR (E309Q) resulted in complete loss of activity, confirming its catalytic function (Figure S5), suggesting that Epi-STR shares a similar mode of substrate binding and catalytic mechanism with the canonical STR enzyme. A five residue-loop near the active site in RsSTR is only present in enzymes from plants in the Apocynaceae family where it was suggested to interact with the glucose moiety of the secoiridoid substrate (e.g. *C. roseus* and *R. serpentina*, but not *O. pumila* and *P. speciosus*) (Figure 2a,c)^[17]^. Docking of vincosidic acid **4R** to the Epi-STR alphafold3 model suggests that the secoiridoid moiety could occupy some of the space where the loop is located in RsSTR (Figure 2d, e, S6).

**Figure 2.**
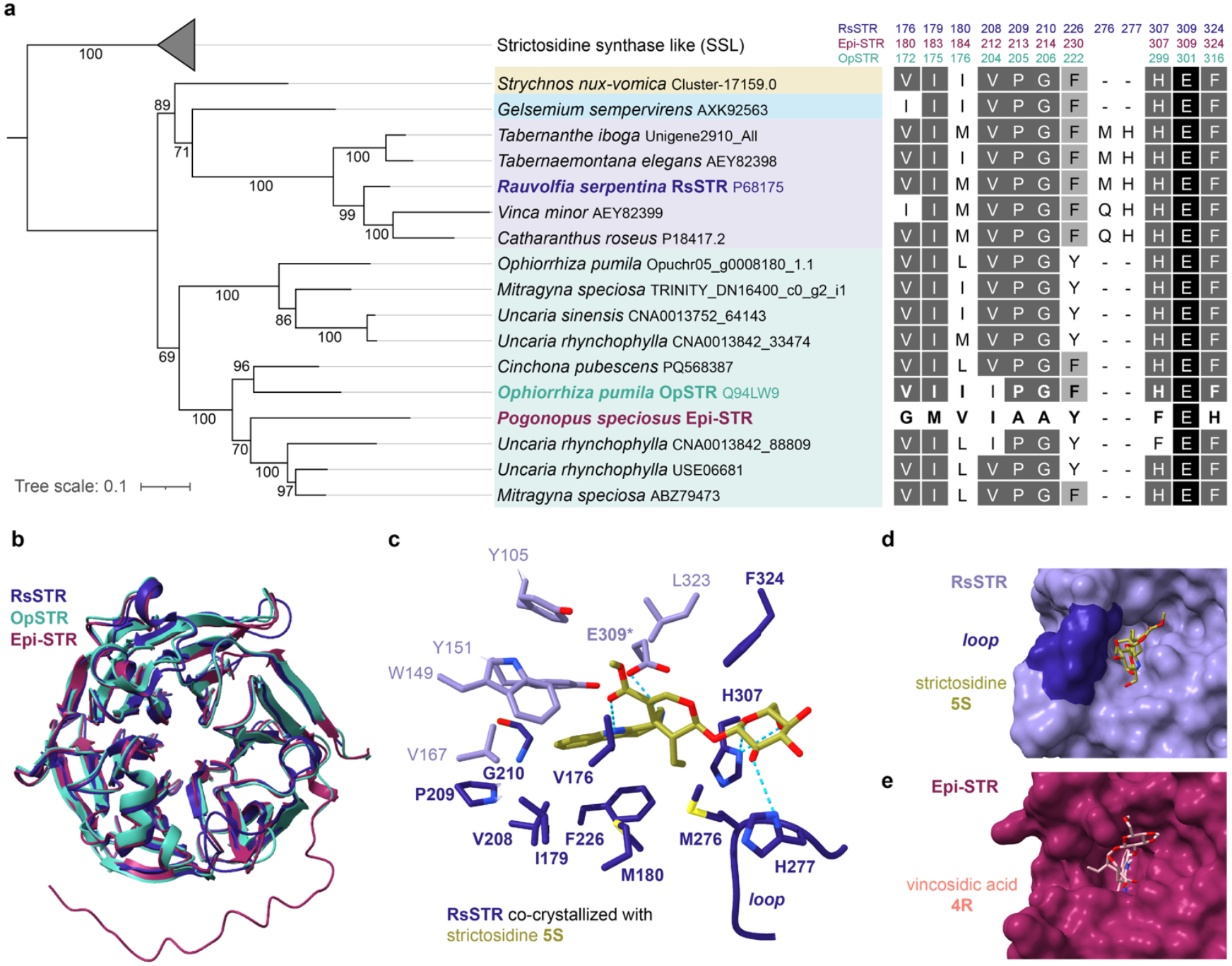
Phylogenetic and structural comparison of Epi-STR with canonical STRs. **a**, Phylogenetic tree of Epi-STR and canonical STR orthologs from MIA-producing Gentianales families: Loganiaceae (beige), Gelsemiaceae (light blue), Apocynaceae (light purple), and Rubiaceae (green) with Epi-STR shown in burgundy. SSLs (strictosidine-synthase-like proteins) ubiquitously occur in plants but do not function as STRs and served as outgroup here. A non-collapsed version of this tree is shown in Figure S4. The alignment on the right depicts active site residues that are substituted or absent in Epi-STR compared to canonical STRs in addition to the strictly conserved catalytic glutamate residue. Amino acid residues targeted for reciprocal substitutions in OpSTR and Epi-STR are depicted in bold. The different shades depict the level of amino acid residue identity among the shown STRs (excluding SSLs): black, identical in 100% of the sequences, dark gray, 80-99%, light gray: 55-79%, white: <55%. Numbers of amino acid residues corresponding to unprocessed sequences including predicted subcellular transit peptides of RsSTR, OpSTR and Epi-STR, respectively, are shown on top. **b**, Overlay of crystal structures of RsSTR (PDB code: 2v91), OpSTR (PDB code: 6s5j) and an Alphafold3 model of Epi-STR ^[7, 16]^. Note, that the C-terminus is unresolved in the crystal structures and therefore only appears in the Epi-STR model. **c**, Depiction of the side chains of residues of the active site of RsSTR co-crystallized with strictosidine (PDB code: 2v91) ^[16]^. Residues that are conserved in Epi-STR are shown in light purple, with the catalytic glutamate in bold and labelled with an asterisk. Residues that are substituted or absent in Epi-STR are shown in dark blue and bold (see alignment in **a**). The loop consists of five residues, only the side chains of the two residues previously shown to be involved in secoriridoid coordination are shown ^[16]^. **d**, Surface view of RsSTR in complex with strictosidine **5S**, the loop is colored in dark (PDB code: 2v91). **e**, Alphafold3 model of Epi-STR with docked vincosidic acid **4R** showing that the secoirdoid moiety would partly occupy the space where the loop is located in Apocynaceae-specific STRs (compare to **d** and S6).

We next aimed to assess how specific active site residue substitutions impacted the stereoselective outcome. For this analysis, we expressed *OpSTR, RsSTR* and *Epi-STR* wild-type and mutant enzymes transiently in *N. benthamiana* and fed tryptamine **1** together with either secologanic acid **2** or secologanin **3** as substrates to leaf disks (Figure 3, S7). All three wild-type orthologs maintained stereoselectivity when either secoiridoid was used as the aldehyde substrate (Figure 3h,i, S7). We observed a striking positional swap of a histidine (H299 in OpSTR; F307 in Epi-STR) and a phenylalanine residue (F316 in OpSTR; H324 in Epi-STR) in Epi-STR (Figure 2a, 3 a, d).

**Figure 3.**
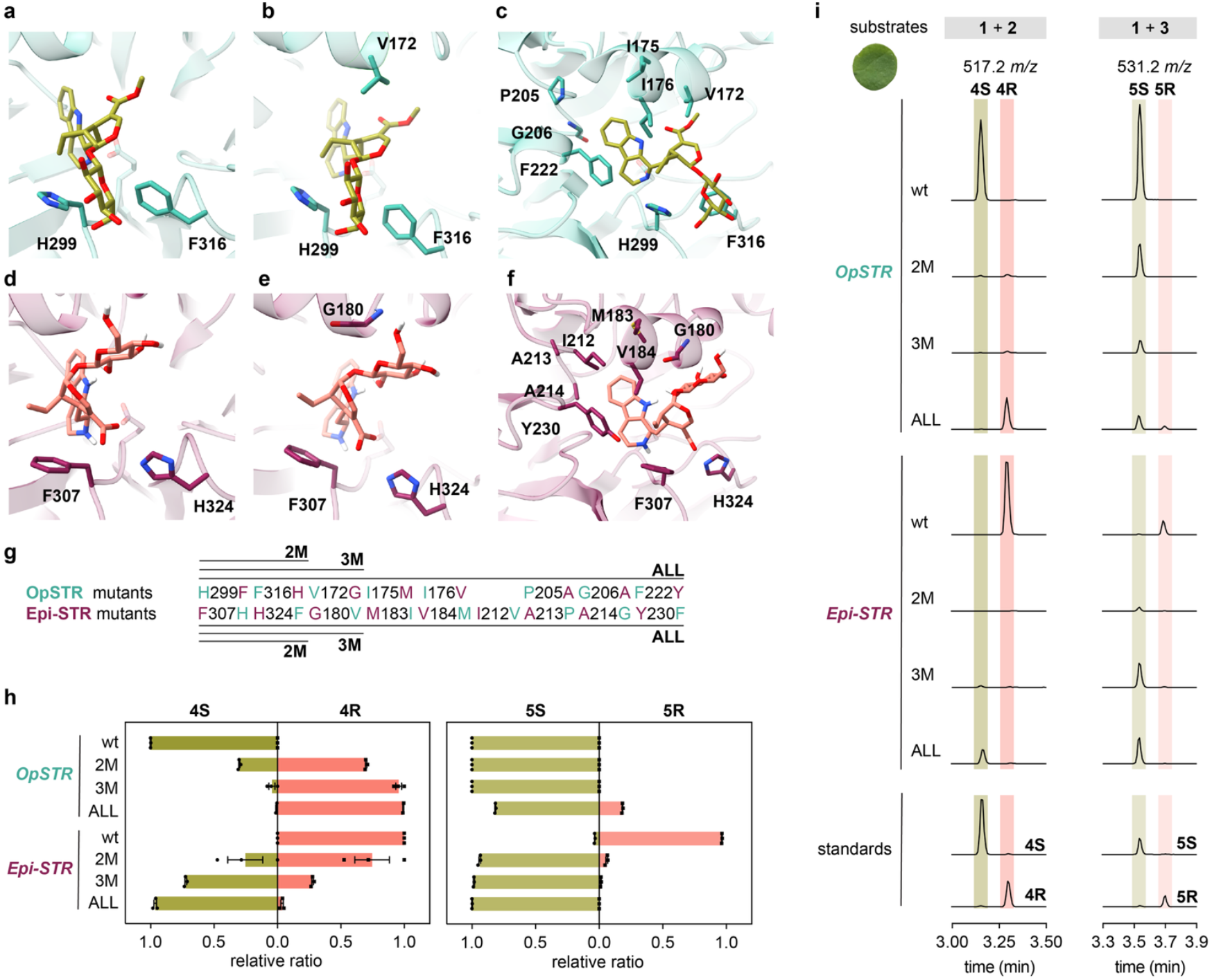
Directing stereopreference through rational reciprocal substitutions in OpSTR and Epi-STR. **a**-**c**, Crystal structure of OpSTR (PDB: 6s5m) with strictosidine **5S** superimposed from RsSTR in complex with strictosidine **5S** (PDB: 2v91). From left to right: residues sequentially targeted for mutation. **d**-**f**, Epi-STR Alphafold3 model with vincosidic acid **4R** docked in active site. From left to right: residues sequentially targeted for mutation. **g**, Mutant details. **h, i**, Expression of *OpSTR* and *Epi-STR* wild-type and mutants in *N. benthamiana* and feeding of tryptamine **1** and secologanic acid **2** or secologanin **3** as substrates to leaf disks. **h**, Relative product ratio of *S* versus *R* epimers. Bar graphs depict the relative ratio of the mean peak area value of three biological replicates of each epimer compared to the combined peak area values of both epimers. Error bars depict standard errors of the mean. Symbols are values of individual biological replicates. **i**, Extracted ion chromatograms (EICs) of one representative replicate from the experiment shown in **h.** Traces in the same column are shown in the same scale to compare product amounts between mutants. Peak area values for all three biological replicates are shown in Figure S7b-c. The results were independently corroborated by in vitro assays with recombinantly produced enzymes (Figure S8).

Both residues are highly conserved among canonical STRs and suggested to coordinate the glucose moiety of the secoiridoid substrate ^[17]^. In a model of Epi-STR, the glucose moiety of vincosidic acid **4R** points in the opposite direction (Figure 3d). Hence, we hypothesized that the swap of this histidine and phenylalanine residues would lead to a different orientation of the secoiridoid substrate and thereby contribute to the observed opposite outcome in stereoselectivity. Indeed, with secologanic acid **2** as a substrate, the OpSTR double mutant (H299F, F316H), predominantly produced vincosidic acid **4R** (Figure 3h). However, the reciprocal Epi-STR mutant (F307H, H324F) maintained stereopreference for vincosidic acid **4R** and only produced strictosidinic acid **4S** as a minor product (Figure 3h). In contrast, with secologanin **3** as substrate, the OpSTR double mutant maintained the native *S*-selectivity whereas the Epi-STR double mutant almost exclusively produced strictosidine **5S** (Figure 3h). Product amounts were reduced for all double mutants compared to wild-type enzymes (Figure 3i). The corresponding RsSTR double mutant (H307F, F324H), largely maintained *S*-preference when secologanic acid **2** was used as the aldehyde substrate (Figure S7a). Since this may be due to the steric hinderance of the Apocynaceae-specific loop and coordination of the glucose moiety by the loop’s histidine residue (H277) (Figure 2c, d, S6), we used OpSTR (Rubiaceae) for further engineering analysis (Figure 3g). Canonical STR enzymes have either a valine or an isoleucine residue near the methylester group of strictosidine **5S** (V172 in OpSTR; Figure 3b)^[16]^. In Epi-STR this hydrophobic residue is substituted with glycine (G180) and located near the glucose of the secoiridoid moiety, leading us to speculate that this residue may be important for stereoselectivity (Figure 3e). Consistent with this hypothesis, reciprocal valine glycine substitutions led to further enhanced changes in stereopreference when secologanic acid **2** was used (Figure 3h). The OpSTR triple mutant (H299F, F316H, V172G) produced vincosidic acid **4R** at a much higher selectivity, and, correspondingly, the Epi-STR triple mutant (F307H, H324F, G180V) predominantly produced strictosidinic acid **4S** (Figure 3h). However, both triple mutants are *S*-selective when secologanin **3** is used as the substrate (Figure 3h). We then introduced reciprocal substitutions in all identified active site residues that differed between orthologs: OpSTR_ALL and Epi-STR_ALL (Figure 2a, 3c, f, g). This more extensive mutagenesis led to nearly complete switches in stereoselectivity with secologanic acid **2** as substrate, and moreover, increased product amounts were observed when compared to triple mutants (Figure 3h, i). When secologanin **3** was used as substrate, the OpSTR_ALL mutant produced modest levels of vincoside **5R** (Figure 3h) but product amounts did not increase compared to triple mutants. Assays of secologanin with Epi-STR_ALL were nearly identical to the triple mutant (F307H, H324F, G180V) (Figure 3 h,i). These results obtained using leaf disk assays were corroborated with in vitro assays using recombinantly produced enzymes (Figure S8). In summary, a single positional swap of two amino acid residues greatly determines stereopreference, but additional active site residue substitutions are needed to fix stereoselectivity and to improve efficiency. Finally, the results indicate that *R*-selectivity is achieved with fewer amino acid mutations when secologanic acid **2** is used as a substrate. It is thus tempting to speculate that the absence of the methylester due to the loss of LAMT activity in *P. speciosus* may have facilitated the evolution of *R*-stereoselectivity.

In conclusion, we discovered the previously unreported accumulation of the MIA vincosidic acid **4R** in *P. speciosus*. The appearance of this unusual stereochemical preference for this Pictet-Spengler product is due to an evolutionary divergent ortholog of a canonical Pictet-Spenglerase, STR. Since MIA biosynthesis is ancestral to this order of plants, we speculate that *P. speciosus* ancestors produced MIAs prior to appearance of the related ipecac alkaloid pathways for which this plant is best known and retained the scaffold-forming *STR* gene ^[3c, 8d, 10]^. *R*-selectivity of this reaction may have emerged to avoid competition with ipecac alkaloid pathway enzymes, which utilize similar substrates. Experiments are underway to unravel the biological functions of vincosidic acid **4R** which may lead to more insights into the reasons why *R* selectivity emerged. This work demonstrates the importance of modern phytochemical analyses of poorly described medicinal plant species, shows how stereoselectivity can be switched by very few specific amino acid substitutions, and provides access to epimer-pure vincosidic acid **4R** and vincoside **5R**.

## Supporting Information

The supporting information contains all experimental details, gene sequences and additional results. The authors have cited additional references within the Supporting Information ^[3c, 8c, 11, 18]^.

## Acknowledgements

We thank Dr. Nils Köster and the Botanical Garden Berlin for generously sharing *P. speciosus* material, and Sabine Scheel, Dr. Klaus Gase and Eva Rothe for help with harvesting. We thank Franz Kaltofen for growing *N. benthamiana* plants, Jens Wurlitzer for assistance with recombinant protein purifications, and Dr. Maritta Kunert for assistance with LC-MS measurements. Dr. Allwin McDonald and Dr. Gyumin Kang are acknowledged for providing previously synthesized authentic standards of vincoside and strictosidine. This work was supported by the Max Planck Society.

