## Supplementary material for "Discovery of a Plant Pictet-Spenglerase with R-Stereoselectivity": Material and Methods; Figures S1-S8; Table S1-S3

**Supporting Information**

### Material and Methods

#### Plant material

*P. speciosus* (IPEN VE-0-B-1680113) was grown at the Botanical Garden Berlin, Germany (Botanischer Garten und Botanisches Museum Berlin). *N. benthamiana* plants for transient gene expression were grown in a greenhouse at 20 - 24 °C, 60 % relative humidity with a 16 h-light / 8 h-dark photoperiod.

#### Sample collection of *P. speciosus* and metabolite extraction

*P. speciosus* samples were collected from the plant grown at Botanical Garden Berlin, snap-frozen in liquid nitrogen and kept at –70°C until further processing. Using pre-chilled mortars and blenders, the plant material was ground to fine powder and stored at −70 °C until extraction.
For metabolite extraction, 10 - 20 mg powder was further ground using metal beads and a TissueLyser II (Qiagen). Extraction was done with 100% MeOH containing 10 mg / L caffeine as internal standard at a ratio of 10 µL MeOH per 1 mg plant sample. Samples were vortexed, sonicated for 10 min in a sonication bath (Bandelin Sonorex) at room temperature, incubated for 15 min at room temperature on a rotator, and centrifuged at 16000 x *g*. The supernatant was further diluted (MeOH / water 1:1), filtered with a 0.22 µm PTFE syringe filter and analyzed by UHPLC-MS/MS method 1.

#### RNA extraction and sequencing

Total RNA was extracted from different *P. speciosus* tissues (Figure S1) using the RNeasy Mini Kit (Qiagen) following the manufacturer's protocol 'Purification of total RNA from Plant Cells and Tissues and Filamentous Fungi' including an on column DNAse digest. To remove phenolic compounds an additional lysis step was included for bark, root and young stem samples as described in the user developed protocol 'Isolation of total RNA from woody plant tissues using the RNeasy Plant Mini Kit' (https://www.qiagen.com/us/resources/resourcedetail?id=9f16addc-8414-4a5d-b969-c620e73b3235&lang=en). This was followed by a 10 min incubation step at 70 °C. Purity and concentration of all samples was measured at a NanoPhotometer® N60 (Implen) and samples were stored at −70°C. 200 ng of total RNA from each tissue were mixed to obtain a pooled RNA sample which was shipped to Novogene for Pacific Biosciences (PacBio) REVIO Sequencing. The following procedures took place at Novogene according to the company's standard procedures: Precise RNA quantity was determined using Qubit and RNA integrity (RIN) value using the Bioanalyzer 2100 system (Agilent technologies). cDNA was synthesized using the Iso-Seq Express 2.0 kit and the library was constructed using the Kinnex Full-Length RNA kit. The library was sequenced on a PacBio Revio SMRT Cell 25M yielding 11 million HiFi reads provided by the company. Processing of HiFi reads was done in-house with IsoSeq 4.0 followed by clustering with CD-HIT (sequence identity threshold 0.98), both run on OmicsBox 3.3 (Biobam). The resulting clustered transcriptome was used to identify Epi-STR.

#### Identification of Epi-STR and Cloning

The amino acid sequence of CrSTR (P18417.2) was blasted against the *P. speciosus* clustered transcriptome assembly. The transcript with the highest sequence identity (50.96%) was cloned as follows. cDNA was reverse transcribed from the RNA (see above) using Superscript VILO™ Master Mix (Thermo Fisher Scientific) in a 20 µL reaction with 2.5 µg RNA input according to manufacturer's instructions and used as template for amplification of Epi-STR (wild-type sequence). Codon-optimized Epi-STR and all other genes and mutants were amplified from synthetic genes (Twist Bioscience). Genes were amplified using Q5 high fidelity polymerase 2x PCR Master Mix (New England Biolabs) according to the manufacturer's protocol and using the primers listed in Table S1. Amplified fragments were gel-purified using the Zymoclean Gel DNA recovery kit (Zymo) according to the manufacturer's instructions and cloned by In-Fusion cloning with 5x In-Fusion Snap Assembly Master Mix (TaKaRa Bio). For transient gene expression in *N. benthamiana*, the sequences were cloned into BsaI-HF_v2 (New England Biolabs) digested 3Ω1 vector containing *Solanum lyopersicum* UBQ10 promoter and terminator[1]. For recombinant protein production genes were cloned into KpnI-HF and HindIII-HF (New England Biolabs) digested pOPINM vector (for fusion to maltose binding protein to enhance protein solubility)[2]. The reactions were transformed into heat-shock competent *Escherichia coli* TOP 10. Plasmids were isolated using Wizard Plus SV Miniprep DNA purification System kit (Promega) according to manufacturer's instructions and sequences were confirmed by Sanger sequencing.

**Recombinant protein production and purification**

Purified plasmids with the coding sequences in a pOPINM vector backbone were used to transform heat shock competent *E. coli* BL21 cells. The strains were grown over night on LB agar plates at 37 °C. Glycerol stocks were made from single colony cultures and stored at −70 °C.

For precultures, *E. coli* BL21 harboring the plasmids for protein overexpression were transferred from glycerol stocks into LB medium with Carbenicillin and were grown over night at 37 °C and 220 rpm shaking. Precultures were transferred into 2xYT medium with Carbenicillin and grown until $OD_{600}$ of 0.6 was reached. For kinetics measurements a 1 L culture of the codon-optimized Epi-STR strain (Epi-STR codon-optimized) was grown, all other strains were grown in 100 mL cultures. For single time point assays, Epi-STR without codon optimization was used. Protein production was induced with 200 µM IPTG and the cultures were grown over night at 18 °C. The cells were pelleted at 4000 x *g* and flash frozen in liquid nitrogen. The thawed pellet was resuspended in lysis buffer A1 (50 mM TRIS-HCl, 50 mM glycine, 5% v/v glycerol, 0.5 M NaCl, 20 mM imidazole, pH 8) with 0.2 g / L lysozyme and 1 tablet / 50 mL buffer of cOmplete™ EDTA-free Protease Inhibitor (Roche) and incubated for 30 min on ice, before lysis by sonification for 2 min (2 s on, 3 s off) on ice (Bandlein UW 2070). The cell debris was pelleted and the supernatant was incubated with Ni-NTA beads (Qiagen) for 1.5 h at 4 °C under gentle shaking. The slurry was pelleted and the beads were washed with wash-buffer (A1) before the proteins were eluted with buffer with high imidazole content (as A1 but with 500 mM imidazole). Elution fractions were concentrated using Amicon Ultra Centrifugal filters (Merck, Germany) by sequential dilution using storage buffer (20 mM 4-(2-hydroxyethyl)-1-piperazineethanesulfonic acid (HEPES) and 150 mM NaCl, pH 7.5). Protein concentrations were determined using the NanoPhotometer® N60 (Implen) by measuring the absorbance at 280 nm and using the calculated extinction coefficient. Purified enzymes were flash-frozen in small aliquots and stored at −70 °C.

**Kinetic measurements with 6xHis_MBP_Epi-STR**

Assays were performed in three technical replicates in 20 µL reaction volumes at 30 °C and 400 rpm shaking. Assays contained 50 mM HEPES pH 7.0 and 1 µM purified Epi-STR (codon optimized, see above). To determine the $K_m$ of secologanic acid, tryptamine concentration was fixed at 1.5 mM and secologanic acid concentration varied between 0.005 and 5 mM. To determine the $K_m$ of tryptamine, secologanic acid concentration was fixed at 2 mM and tryptamine concentration varied between 0.1 and 2000 µM. For $K_m$ of secologanin, a fixed tryptamine concentration 1.5 mM was used and secologanin concentration varied between 0.005 and 5 mM. Time point samples were taken every 15 min by flash-freezing in liquid nitrogen. For LC-MS measurements samples were diluted 1:50 in 70% MeOH (+ caffeine 0.5 mg/L as internal standard) immediately prior to UHPLC-MS measurement. Only for the reaction with secologanin, 0.1% formic acid was added to the 70% MeOH to prevent the spontaneous lactamization of vincoside. Samples were filtered through a 0.45 µm low-binding PTFE filter plate (MultiScreen® Solvinert 96, Merck-Millipore) into a 96-well microtiter plate (SureSTART™ WebSeal™, Thermo Scientific) according to the manufacturer's instructions. Plates were sealed with Rapid Slit Seal (BioChromato) and analyzed with UHPLC-MS method 2.

**Single time-point assays**

Assays were performed in three technical replicates and contained 50 mM HEPES, 1 µM enzyme and 1 mM of tryptamine **1** and secologanic acid **2** or secologanin **3**, respectively, in a 50 µL total assay volume. Assays were incubated at 30°C and 400 rpm shaking and stopped after 3.5 h by flash-freezing in liquid nitrogen. The samples were diluted 1:20 in 70% MeOH containing 0.1% formic acid and 0.5 mg/L caffeine as internal standard. Samples were filtered through a plate as described above and analyzed with UHPLC-MS method 2.

**_A. tumefaciens_ mediated transient gene expression in _N. benthamiana_**

*Agrobacterium tumefaciens GV3101* was transformed with the respective plasmids through electroporation. After transformation, the cells were recovered in YEB medium without antibiotics and incubated on YEB agar plates with rifampicin, gentamycin, and spectinomycin at 28 °C for two days. For glycerol stock preparation, single colonies were picked, confirmed by colony PCR and grown in YEB medium containing antibiotics for 24 h at 28 °C. Stocks were stored at −70 °C. The stocks were used for agroinfiltration as follows [3]. Aliquots of the glycerol stocks were spread on YEB agar plates containing antibiotics and 100 µM acetosyringone. The cells were grown over night at 28 °C and gently resuspended in 1-2 mL infiltration buffer (10 mM 2-(N-morpholino)ethanesulfonic acid (MES) pH 5.7; 10 mM $MgCl_2$; 100 µM acetosyringone). The $OD_{600}$ was measured using and OD600 DiluPhotometer™ (Implen). The strains were mixed and diluted with infiltration buffer to $OD_{600}$=0.1 per strain. A strain harboring the plasmid for overexpression of the RNA silencing suppressor *P19* was co-infiltrated in all cases. The mixtures were infiltrated into the leaves of 3-4 weeks old *N. benthamiana* plants (1 leaf per plant). Plants were grown under grow lights (16 h/8 h light/dark) to allow expression of transgenes. Three days after agroinfiltration 1 cm leaf disks (1 disk per leaf) were cut using a leaf puncher and incubated in 200 µL of 25 mM HEPES buffer pH 7 containing 200 µM of each substrate (as indicated) in 48-well plates. Plates were sealed with parafilm and incubated for 24 h under grow lights (16 h/8 h light/dark).

**Metabolite extraction from _N. benthamiana_ leaf disks**

Each leaf disk was transferred into a 2 mL Eppendorf tube containing 1 metal bead, flash frozen in liquid nitrogen, and ground using a TissueLyser II (Quiagen) with precooled adaptors. The samples were then extracted with 100 µL of 70% MeOH containing 0.1% formic acid and 1 mg/L caffeine. The samples were vortexed, sonicated for 10 min (Bandelin Sonorex), incubated on a rotator for 15 min and centrifuged at 16 000 x *g*. The supernatants were filtered through a 0.45-µm low-binding PTFE filter plate (MultiScreen® Solvinert 96, Merck-Millipore) into a 96-well microtiter plate (SureSTART™ WebSeal™, Thermo Scientific) according to the manufacturer's instructions. Plates were sealed with Rapid Slit Seal (BioChromato) and analyzed with UHPLC-MS/MS method 1.

**UHPLC-MS methods**

Chromatographic separation was performed on an UltiMate 3000 Ultra High-Performance Liquid Chromatography (UHPLC) system (Thermo Fisher Scientific) equipped with a Kinetex XB-C18 (2.1 x 100 mm, 2.6 µm; 100 Å) column (Phenomenex) with an oven temperature of 40 °C. The flow rate was set to 0.6 mL/min and the injection volume was 2 µL. The mobile phase consisted of water with 0.1% formic acid (solvent A) and acetonitrile (solvent B) and the gradient was as follows: 5% B at 0-1min, to 50% B at 6 min, to 100% B at 6.1 min. After that, the column was flushed with 100% B until 7.5 min and re-equilibrated to 5% B until 10 min. The UHPLC system was coupled to an Impact II high resolution Quadrupole Time-Of-Flight (QTOF) mass spectrometer (Bruker Daltonics, Massachusetts, USA) with an electrospray ionization (ESI) operated in positive ionization mode. The source parameters were set as follows: 3500 V capillary voltage, 500 V end plate offset; nebulizer pressure of 2.5 bar, with nitrogen at 250 °C and a flow of 11 L/min as the drying gas. At the first minute of each run the LC input was redirected to waste. During this time, the m/z values of the instrument were calibrated using the cluster ion *m/z* values of a sodium formate-isopropanol solution injected by direct source infusion with a 5 mL syringe connected to an external pump at a flow rate of 0.18 mL/h.

Two different acquisition methods were used: Method 1 involved data-dependent MS/MS detection to obtain fragment spectra of specific precursors. These were recorded at a frequency of 12 Hz and a mass range of 80–1000 m/z with data-dependent MS/MS and an active exclusion window of 0.2 min, a review threshold of 1.8 fold. Fragmentation was triggered on an absolute threshold of 400 counts with a total cycle time limited to 0.5 s. The step model (from 20 to 50 eV) was used for the collision energy. Method 2 consisted of data acquisition exclusively in full scan mode. For this purpose, the spectral rate was set to 2 Hz and a mass range of 80 to 1000 m/z was selected.

**MS data analysis**

Data was analyzed with MZmine 3.6.0-4.5.0 [4] and DataAnalysis 5.0 (Bruker). Traces of extracted ion chromatograms (EICs) were exported from MZmine. Peak areas were determined using the MZmine processing wizard. Peak areas were normalized to the internal standard caffeine and converted into intensity per second. For kinetics analysis product concentrations were calculated in Microsoft Excel 2021 using standard curves. Further analysis and construction of graphs was done in GraphPad Prism 10. For kinetic analysis, concentrations of vincosidic acid **4R** or vincoside **5R**, respectively, were calculated by linear regression against a standard concentration curve GraphPad Prism 10. Kinetic parameters were calculated using the Michaelis-Menten equation in GraphPad Prism 10.

**Chemicals used in this study**

Secologanin (50741) and tryptamine hydrochloride (246557) were purchased from Sigma. Strictosidine **5S** and vincoside **5**R authentic standards were obtained from previously synthesized stocks[5]. Secologanic acid **2** was synthesized by alkaline hydrolysis of secologanin **3** as previously described [3a, 6]. Briefly, secologanin **3** was incubated in 0.1 M NaOH at room temperature for 5 h (40 µL per 1 mg secologanin). The reaction was stopped by neutralization using HCl (10 M stock) and completeness of the reaction was confirmed by LC-MS analysis. Vincosidic acid **4R** and strictosidinic acid **4S** were obtained through enzymatic de-esterification of vincoside **5R** or strictosidine **5S**, respectively, using previously purified recombinant *Carapichea ipecacuanha* deacetyl(iso)ipecoside esterase (CiDE)[3a]. The reaction mixtures (50 µL total volume) contained 1 mM CiDE, 0.5 M HEPES (pH 7.0) and 0.1 mM of vincoside **5R** or strictosidine **5S**. Reactions were incubated for 16h at 30°C and 600 rpm shaking. Reactions were stopped by adding 50 µL of 100% MeOH and the completeness of the reactions was confirmed by UHPLC-MS/MS method 2.

**Protein models and docking**

The structural model of Epi-STR was predicted using the AlphaFold 3 server (https://alphafoldserver.com/)[7]. Numbers of crystal structures obtained from the PDB database (https://www.rcsb.org/) are mentioned in each figure legend. Docking studies were performed with AutoDock Vina on the SwissDock webserver (https://www.swissdock.ch)[8]. Protein structures and structure models were visualized using ChimeraX 1.8[9]. Chemical structures were drawn in ChemDraw Professional 23.1.2.

**Phylogenetic analyses**

For the generation of the phylogenetic tree, sequences were obtained through BLAST searches against publicly available databases (NCBI and 1KP) and previously published transcriptome assemblies using Epi-STR and CrSTR (P18417.2) amino acid sequences as baits[1, 10]. Table S3 lists all accession numbers and amino acid sequences. Full-length amino acid sequences were aligned using WebPRANK (https://www.ebi.ac.uk/goldman-srv/webprank/)[11]. A maximum-likelihood tree was generated using the IQ-TREE web-server (http://iqtree.cibiv.univie.ac.at/) with automatic substitution model and maximum number of bootstrap replicates of 1000[12]. iTOL (https://itol.embl.de) and Adobe Illustrator 2025 were used to visualize and graphically edit the tree[13].

### Supplementary Figures

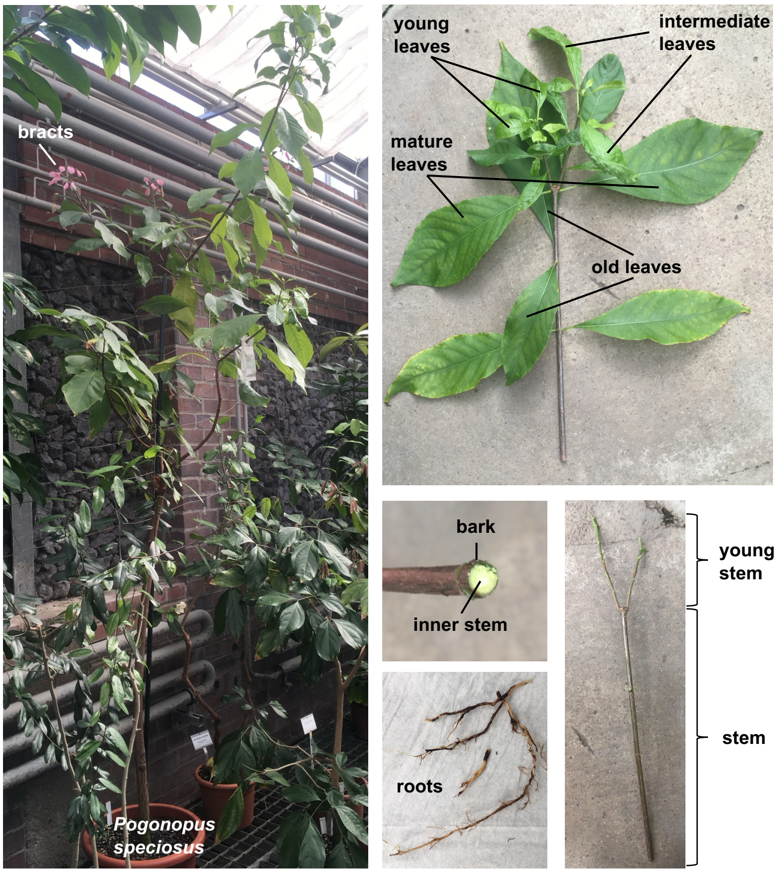

**Figure S1. Photo of *Pogonopus speciosus* plant sampled for RNA-seq.** Plant grew at Botanical Garden Berlin, Germany (see methods). RNA was extracted from indicated tissues as described in the methods. Metabolite analysis shown in this study was done on young leaves sample.

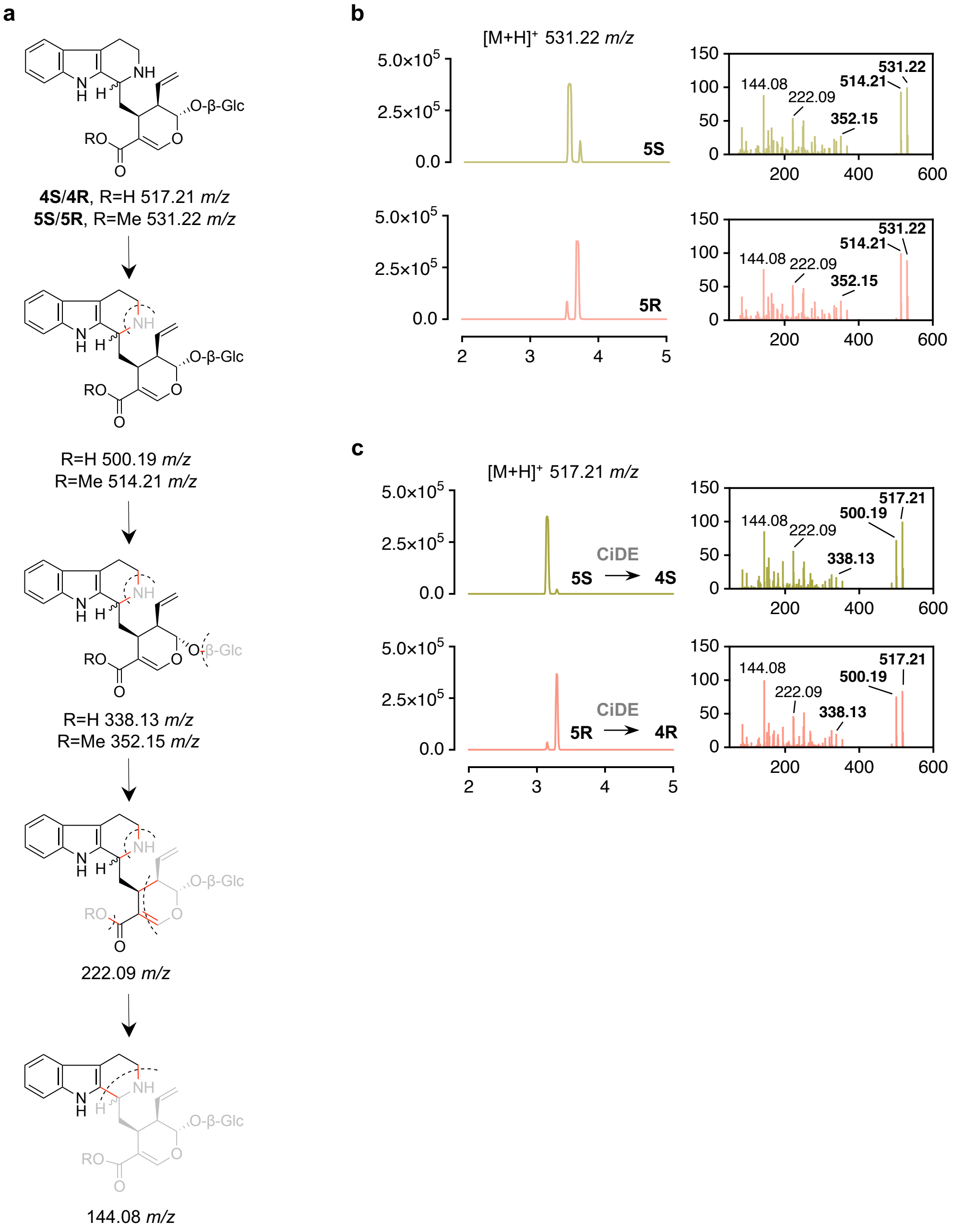

**Figure S2. Extracted ion chromatograms and MS$^2$ data of product standards used in this study.** **a**, Proposed fragmentation of strictosidine **5S**, vincoside **5R,** strictosidinic acid **4S**, and vincosidic acid **4R**. **b-c**, Left, extracted ion chromatograms. Right, corresponding MS$^2$ data shown as relative abundance of ions with *m/z* values of proposed fragment ions indicated. Fragments showing expected differences between methylesters and acids are in bold. **b**, **5S** and **5R** obtained from previously characterized stocks [5]. **c**, Epimer-pure **4S** and **4R** standards were produced through enzymatic de-esterification of **5S** or **5R**, respectively, using an excess of recombinant *Carapichea ipecacuanha* deacetyl(iso)ipecoside esterase (CiDE) [3a] (see methods).

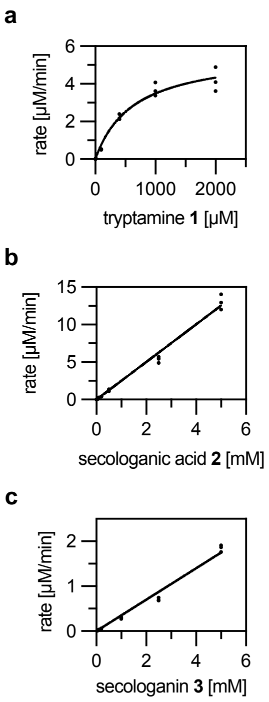

**Figure S3. Steady-state kinetics of Epi-STR.** In vitro assays with purified recombinant codon-optimized Epi-STR. **a**, catalytic activity for tryptamine **1** at 2 mM secologanic acid **2**. **b**, catalytic activity for secologanic acid **2** at 1.5 mM tryptamine. **c**, catalytic activity for secologanin **3** at 1.5 mM tryptamine.

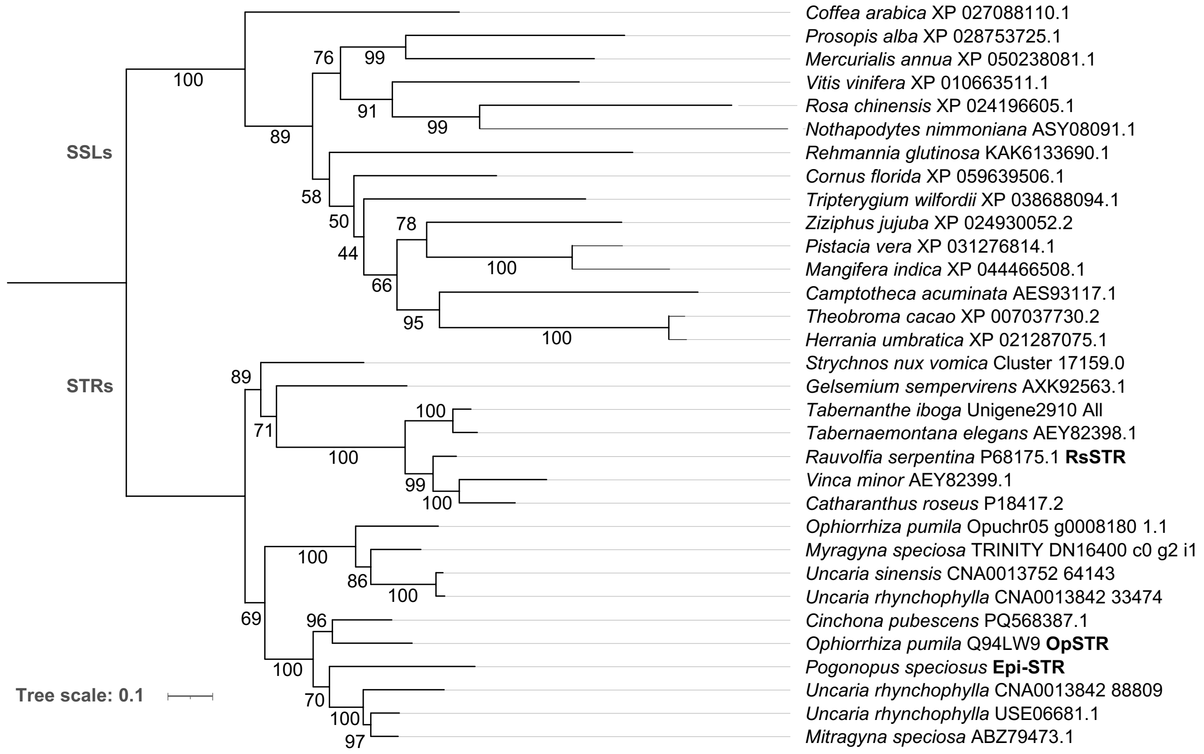

**Figure S4. Maximum likelihood tree of STR amino acid sequences.** Tree from Figure 2a is shown here with non-collapsed strictosidine synthase like sequences (SSLs).

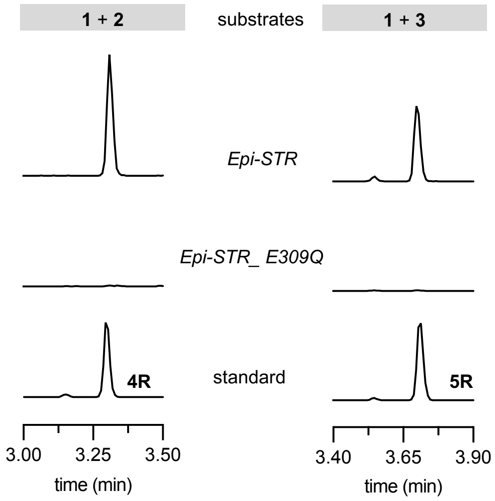

**Figure S5. Mutation of the catalytic glutamate residue in Epi-STR leads to complete loss in activity.** Expression of *Epi-STR* or *Epi-STR_E309Q* mutant in *Nicotiana benthamiana* and feeding with tryptamine **1** and secologanic acid **2** or secologanin **3**. Chromatograms in the same column are shown in the same scale.

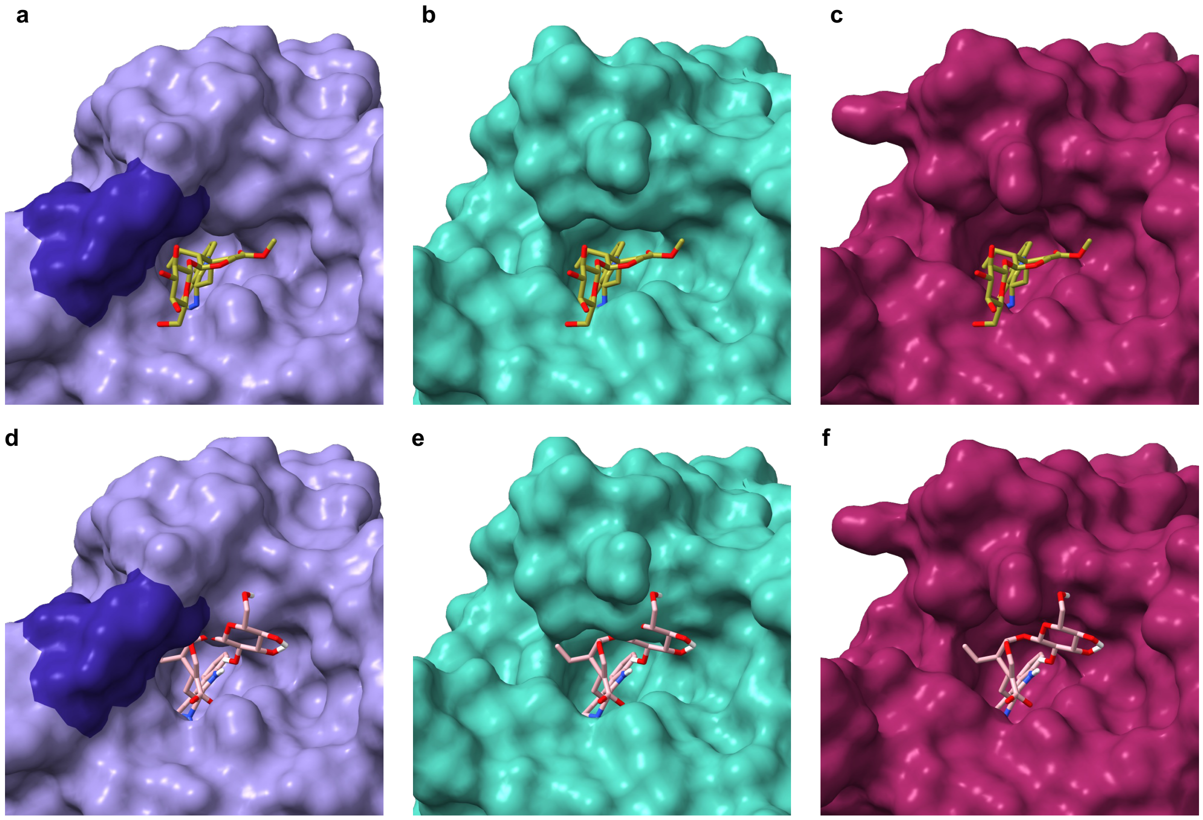

**Figure S6. Surface models of STR active sites with superimposed strictosidine 5S and vincosidic acid 4R.** **a**, RsSTR crystal structure in complex with strictosidine **5S** (PDB code: 2v91) with Apocynaceae-specific loop shown in dark. **b**, OpSTR crystal structure (PDB code: 6s5j) with **5S** superimposed from **a**. **c**, Epi-STR Alphafold 3 model with **5S** superimposed from **a**. **d**, RsSTR crystal structure from **a** superimposed with vincosidic acid **4R** docking from **f**. **e**, OpSTR crystal structure from **b** superimposed with **4R** from **f**. **f**, Epi-STR alphafold 3 model with docked **4R**.

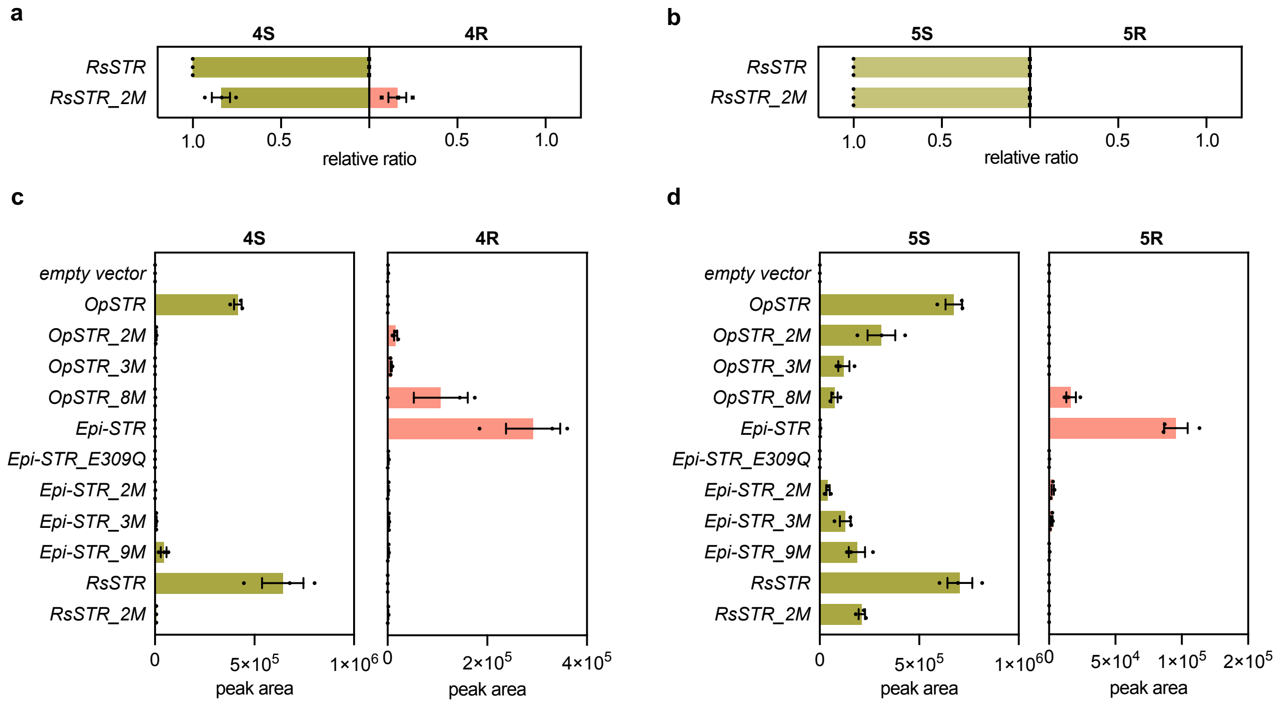

**Figure S7. Expression of *STR* wild-type and mutants in *N. benthamiana*.** Peak area data for the experiment shown in Figure 3 in addition to data for expression of *RsSTR* and *RsSTR_2M*. Genes were expressed in *N. benthamiana* and tryptamine **1** and secologanic acid **2** or secologanin **3** were fed to leaf disks. **a-b**, Relative product ratios of *S* versus *R* epimer. Bar graphs depict the relative ratio of the mean peak area value of three biological replicates of each epimer compared to the combined peak area values of both epimers. Error bars depict standard errors of the mean. Symbols are values of individual biological replicates. See Figure 3h for ratios for other mutants. **a**, Feeding of **1** and **2**. **b**, Feeding of **1** and **3**. **c-d**, LC-MS peak area values shown as bars of the mean of three biological replicates, error bars are standard error of the mean. Symbols depict peak area values of individual biological replicates **c**, Feeding of **1** and **2**. **d**, Feeding of **1** and **3**.

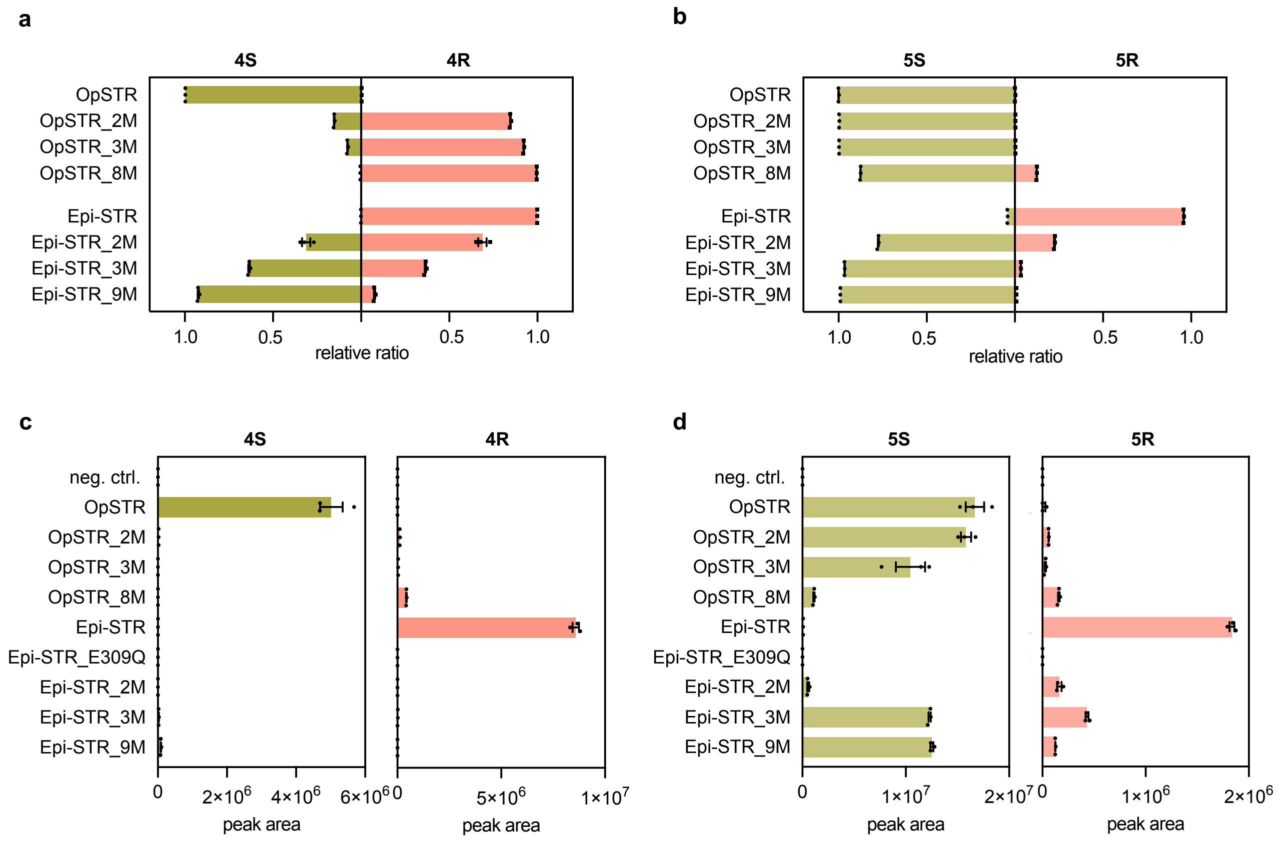

**Figure S8. *In vitro* assays with STR wild-type and mutants.** Purified recombinant STR wild-type and mutants were assayed with 1 mM of tryptamine **1** and secologanic acid **2** or secologanin **3**, respectively, as substrates. Assays were run for 3.5 hours and products were analyzed by LC-MS. **a-b**, Relative product ratios of *S* versus *R* epimer. Bar graphs depict the relative ratio of the mean peak area value of three technical replicates of each epimer compared to the combined peak area values of both epimers. Error bars depict standard errors of the mean. Symbols are values of individual replicates. **a**, Reaction with **1** and **2**. **b**, reaction with **1** and **3**. **c-d**, LC-MS peak area values from the same experiment, shown as bars of the mean of three technical replicates, error bars are standard error of the mean, symbols are peak area values for individual replicates. **c**, Reaction with **1** and **2**. **d**, Reaction with **1** and **3**.

### Supplementary Tables

**Table S1. List of primers used in this study for wild-type and mutant sequences.**

| Gene name | sequence | vector |
|---|---|---|
| Epi-STR | TTTATGAATTTTGCAGCTCGATGCGTAGTTCTGAAAGTATGCT | 3Ω1 |
| | GACAACCACAACAAGCACCGTCAGGCAGAAGCAGACACTC | |
| OpSTR | TTTATGAATTTTGCAGCTCGATGCATAGTTCAGAAGCCATGG | |
| | GACAACCACAACAAGCACCGTTAGAAAGAAGAAAATTCCTTGAATG | |
| RsSTR | TTTATGAATTTTGCAGCTCGATGGCCAAACTTTCTGATTCGC | |
| | GACAACCACAACAAGCACCGTTAATGACTTGAAACAAAAGAA | |
| Epi-STR_co (codon optimized) | AAGTTCTGTTTCAGGGCCCGTCGCCGCTTTTTTTCCAGTT | pOPINM |
| | ATGGTCTAGAAAGCTTTATGCTGATGCACTTACTCCAT | |
| Epi-STR | AAGTTCTGTTTCAGGGCCCGTCGCCGCTGTTTTTCCAGTT | |
| | ATGGTCTAGAAAGCTTTAGGCAGAAGCAGACACTCCAT | |
| OpSTR | AAGTTCTGTTTCAGGGCCCGCATAGTTCAGAAGCCATGGTTGT | |
| | ATGGTCTAGAAAGCTTTAGAAAGAAGAAAATTCCTTGAATG | |

**Table S2. Coding sequences used in this study.**

| Name | Sequence |
|---|---|
| *Epi-STR* | ATGCGTAGTTCTGAAAGTATGCTTGCAGTAAGCATTTTCTTGGTCCTTTTCTCG<br>TCCTCCCCTTCGCTTGTTCTATCTTCGCCGCTGTTTTTCCAGTTTCTTAAGTCA<br>CCCTTTGCCCCCAATGCCTACGGTTTTGGTGCATCTGGTGAACTTTACGCCTC<br>CGTCGAAGATGGCAGAATTGTTAAGTATGAAGGATCGAGCAACAGGTTTTTGA<br>CCCACGCTGTTGCCTCTCCGCACTGGAAGAGAGATCTGTGTGAGAACAACACT<br>AATCCTCAGCTAAAACCCTTCTGTGGGAGAATATATGACTTCGGATTCAACTAT<br>GGAACTCAGCAATTGTACATTGCTGATTCTTATTACGGTCTTTGTGTGGTTGGA<br>GCTAAAGGAGGCCGTGCAACTCAACTTGTCACGAGTGCAGATGGAGTGGACT<br>TTAAGTGGCTTTATGGCTTAGCTTTGGACCAGCATACTGGCATTGTTTACTTCA<br>CAGATGTTAGCTCAGTGTATGATGACAGAGGTGGTCAAGATATGGTGAGGACA<br>AAAGATAAAACAGGAAGATTACTCAAATACGATCCCTCAACAGATGAAGTCACA<br>GTTTTGATGAAAGAGCTGGGTATAGCCGCCGGTACCGAAGTCAGCGAAGATG<br>GCTCGTTTGTTGTTGTTGCTGAATACTTGAGCAATACAATTTTCAAGTATTGGTT<br>AAAGGGTCCGAAGGCAAACACTTATGAAGCCTTACTGACAATCGGGGGCCCA<br>GGGAACATCAAGAGGACTAAAGCTGGAGATTATTGGGGAGCCTCAAGTGATAA<br>TGTTGGAACCAAAGTTATCACAAGCGGTATAAGGTTTGATGAATCTGGCAAAAT<br>TTTAGAAGTCGTGCGTATCCCTCCACCGTATCAAGGCGAATTTTTCGAACAAGT<br>TCATGAGCATAATGGGGCACTTTATATCGGGACTTTGCACCACGACTTTGTAG<br>GTATATTGCACAATTACCAGGGTGCATCTGATCTGAAGGGAAATAATCTAGCTG<br>GGGCCAATGGAGCTTTGAATGGAGTGTCTGCTTCTGCCTGA |
| *Epi-STR_codon optimized* | TCGCCGCTTTTTTTCCAGTTTCTCAAGTCGCCATTTGCGCCGAACGCCTACGG<br>TTTCGGGGCGAGTGGTGAACTGTACGCGTCAGTTGAGGATGGCCGAATTGTG<br>AAGTATGAGGGTAGTTCTAATAGATTTCTTACCCATGCAGTGGCGTCCCCGCA<br>TTGGAAGCGAGATCTGTGTGAGAACAACACCAACCCCCAATTAAAACCGTTTT<br>GCGGACGGATATATGACTTCGGGTTTAATTACGGAACTCAGCAGTTATATATC<br>GCTGACTCATATTATGGCTTATGTGTTGTAGGTGCGAAAGGTGGTCGTGCGAC<br>CCAACTGGTCACGAGCGCAGATGGTGTCGACTTTAAATGGCTGTACGGACTG<br>GCTTTGGACCAGCATACCGGTATTGTCTATTTTACCGATGTTTCTAGCGTGTAT<br>GATGATCGTGGCGGTCAAGACATGGTACGCACAAAAGATAAAACCGGGCGTTT<br>GTTAAAGTATGATCCATCAACAGATGAAGTTACAGTTTTAATGAAAGAACTGGG<br>CATTGCTGCAGGAACAGAAGTAAGTGAAGACGGCAGTTTCGTGGTGGTTGCT<br>GAATATTTGTCAAATACTATCTTTAAGTATTGGTTGAAAGGTCCAAAAGCTAATA<br>CGTATGAAGCATTGCTCACAATTGGCGGTCCTGGAAACATTAAGCGTACAAAA<br>GCTGGAGATTATTGGGGAGCATCATCTGACAATGTCGGTACAAAAGTCATTAC<br>ATCCGGCATCCGCTTCGACGAGAGTGGGAAGATTTTGGAGGTTGTGCGAATTC<br>CGCCCCCTTACCAGGGCGAATTCTTCGAACAAGTTCACGAACACAACGGAGCT<br>TTATATATTGGCACGTTGCATCATGATTTCGTTGGAATTCTGCACAATTACCAG<br>GGAGCGAGCGATCTGAAAGGAAACAACCTGGCAGGAGCGAATGGAGCGCTTA<br>ATGGAGTAAGTGCATCAGCA |
| *Epi-STR_E309Q* | ATGCGTAGTTCTGAAAGTATGCTTGCAGTAAGCATTTTCTTGGTCCTTTTCTCG<br>TCCTCCCCTTCGCTTGTTCTATCTTCGCCGCTGTTTTTCCAGTTTCTTAAGTCA<br>CCCTTTGCCCCCAATGCCTACGGTTTTGGTGCATCTGGTGAACTTTACGCCTC<br>CGTCGAAGATGGCAGAATTGTTAAGTATGAAGGATCGAGCAACAGGTTTTTGA<br>CCCACGCTGTTGCCTCTCCGCACTGGAAGAGAGATCTGTGTGAGAACAACACT<br>AATCCTCAGCTAAAACCCTTCTGTGGGAGAATATATGACTTCGGATTCAACTAT<br>GGAACTCAGCAATTGTACATTGCTGATTCTTATTACGGTCTTTGTGTGGTTGGA<br>GCTAAAGGAGGCCGTGCAACTCAACTTGTCACGAGTGCAGATGGAGTGGACT<br>TTAAGTGGCTTTATGGCTTAGCTTTGGACCAGCATACTGGCATTGTTTACTTCA<br>CAGATGTTAGCTCAGTGTATGATGACAGAGGTGGTCAAGATATGGTGAGGACA<br>AAAGATAAAACAGGAAGATTACTCAAATACGATCCCTCAACAGATGAAGTCACA<br>GTTTTGATGAAAGAGCTGGGTATAGCCGCCGGTACCGAAGTCAGCGAAGATG<br>GCTCGTTTGTTGTTGTTGCTGAATACTTGAGCAATACAATTTTCAAGTATTGGTT<br>AAAGGGTCCGAAGGCAAACACTTATGAAGCCTTACTGACAATCGGGGGCCCA<br>GGGAACATCAAGAGGACTAAAGCTGGAGATTATTGGGGAGCCTCAAGTGATAA<br>TGTTGGAACCAAAGTTATCACAAGCGGTATAAGGTTTGATGAATCTGGCAAAAT<br>TTTAGAAGTCGTGCGTATCCCTCCACCGTATCAAGGCGAATTTTTCCAACAAGT<br>TCATGAGCATAATGGGGCACTTTATATCGGGACTTTGCACCACGACTTTGTAG |

| | |
|---|---|
| | GTATATTGCACAATTACCAGGGTGCATCTGATCTGAAGGGAAATAATCTAGCTG<br>GGGCCAATGGAGCTTTGAATGGAGTGTCTGCTTCTGCCTGA |
| *Epi-STR_2M* | ATGCGTAGTTCTGAAAGTATGCTTGCAGTAAGCATTTTCTTGGTCCTTTTCTCG<br>TCCTCCCCTTCGCTTGTTCTATCTTCGCCGCTGTTTTTCCAGTTTCTTAAGTCA<br>CCCTTTGCCCCCAATGCCTACGGTTTTGGTGCATCTGGTGAACTTTACGCCTC<br>CGTCGAAGATGGCAGAATTGTTAAGTATGAAGGATCGAGCAACAGGTTTTTGA<br>CCCACGCTGTTGCCTCTCCGCACTGGAAGAGAGATCTGTGTGAGAACAACACT<br>AATCCTCAGCTAAAACCCTTCTGTGGGAGAATATATGACTTCGGATTCAACTAT<br>GGAACTCAGCAATTGTACATTGCTGATTCTTATTACGGTCTTTGTGTGGTTGGA<br>GCTAAAGGAGGCCGTGCAACTCAACTTGTCACGAGTGCAGATGGAGTGGACT<br>TTAAGTGGCTTTATGGCTTAGCTTTGGACCAGCATACTGGCATTGTTTACTTCA<br>CAGATGTTAGCTCAGTGTATGATGACAGAGGTGGTCAAGATATGGTGAGGACA<br>AAAGATAAAACAGGAAGATTACTCAAATACGATCCCTCAACAGATGAAGTCACA<br>GTTTTGATGAAAGAGCTGGGTATAGCCGCCGGTACCGAAGTCAGCGAAGATG<br>GCTCGTTTGTTGTTGTTGCTGAATACTTGAGCAATACAATTTTCAAGTATTGGTT<br>AAAGGGTCCGAAGGCAAACACTTATGAAGCCTTACTGACAATCGGGGGCCCA<br>GGGAACATCAAGAGGACTAAAGCTGGAGATTATTGGGGAGCCTCAAGTGATAA<br>TGTTGGAACCAAAGTTATCACAAGCGGTATAAGGTTTGATGAATCTGGCAAAAT<br>TTTAGAAGTCGTGCGTATCCCTCCACCGTATCAAGGCGAACATTTCGAACAAG<br>TTCATGAGCATAATGGGGCACTTTATATCGGGACTTTGTTTCACGACTTTGTAG<br>GTATATTGCACAATTACCAGGGTGCATCTGATCTGAAGGGAAATAATCTAGCTG<br>GGGCCAATGGAGCTTTGAATGGAGTGTCTGCTTCTGCCTGA |
| *Epi-STR_3M* | ATGCGTAGTTCTGAAAGTATGCTTGCAGTAAGCATTTTCTTGGTCCTTTTCTCG<br>TCCTCCCCTTCGCTTGTTCTATCTTCGCCGCTGTTTTTCCAGTTTCTTAAGTCA<br>CCCTTTGCCCCCAATGCCTACGGTTTTGGTGCATCTGGTGAACTTTACGCCTC<br>CGTCGAAGATGGCAGAATTGTTAAGTATGAAGGATCGAGCAACAGGTTTTTGA<br>CCCACGCTGTTGCCTCTCCGCACTGGAAGAGAGATCTGTGTGAGAACAACACT<br>AATCCTCAGCTAAAACCCTTCTGTGGGAGAATATATGACTTCGGATTCAACTAT<br>GGAACTCAGCAATTGTACATTGCTGATTCTTATTACGGTCTTTGTGTGGTTGGA<br>GCTAAAGGAGGCCGTGCAACTCAACTTGTCACGAGTGCAGATGGAGTGGACT<br>TTAAGTGGCTTTATGGCTTAGCTTTGGACCAGCATACTGGCATTGTTTACTTCA<br>CAGATGTTAGCTCAGTGTATGATGACAGAGGTGTTCAAGATATGGTGAGGACA<br>AAAGATAAAACAGGAAGATTACTCAAATACGATCCCTCAACAGATGAAGTCACA<br>GTTTTGATGAAAGAGCTGGGTATAGCCGCCGGTACCGAAGTCAGCGAAGATG<br>GCTCGTTTGTTGTTGTTGCTGAATACTTGAGCAATACAATTTTCAAGTATTGGTT<br>AAAGGGTCCGAAGGCAAACACTTATGAAGCCTTACTGACAATCGGGGGCCCA<br>GGGAACATCAAGAGGACTAAAGCTGGAGATTATTGGGGAGCCTCAAGTGATAA<br>TGTTGGAACCAAAGTTATCACAAGCGGTATAAGGTTTGATGAATCTGGCAAAAT<br>TTTAGAAGTCGTGCGTATCCCTCCACCGTATCAAGGCGAACATTTCGAACAAG<br>TTCATGAGCATAATGGGGCACTTTATATCGGGACTTTGTTTCACGACTTTGTAG<br>GTATATTGCACAATTACCAGGGTGCATCTGATCTGAAGGGAAATAATCTAGCTG<br>GGGCCAATGGAGCTTTGAATGGAGTGTCTGCTTCTGCCTGA |
| *Epi-STR_9M* | ATGCGTAGTTCTGAAAGTATGCTTGCAGTAAGCATTTTCTTGGTCCTTTTCTCG<br>TCCTCCCCTTCGCTTGTTCTATCTTCGCCGCTGTTTTTCCAGTTTCTTAAGTCA<br>CCCTTTGCCCCCAATGCCTACGGTTTTGGTGCATCTGGTGAACTTTACGCCTC<br>CGTCGAAGATGGCAGAATTGTTAAGTATGAAGGATCGAGCAACAGGTTTTTGA<br>CCCACGCTGTTGCCTCTCCGCACTGGAAGAGAGATCTGTGTGAGAACAACACT<br>AATCCTCAGCTAAAACCCTTCTGTGGGAGAATATATGACTTCGGATTCAACTAT<br>GGAACTCAGCAATTGTACATTGCTGATTCTTATTACGGTCTTTGTGTGGTTGGA<br>GCTAAAGGAGGCCGTGCAACTCAACTTGTCACGAGTGCAGATGGAGTGGACT<br>TTAAGTGGCTTTATGGCTTAGCTTTGGACCAGCATACTGGCATTGTTTACTTCA<br>CAGATGTTAGCTCAGTGTATGATGACAGAGGTGTTCAAGATATAATGAGGACA<br>AAAGATAAAACAGGAAGATTACTCAAATACGATCCCTCAACAGATGAAGTCACA<br>GTTTTGATGAAAGAGCTGGGTGTACCCGGCGGTACCGAAGTCAGCGAAGATG<br>GCTCGTTTGTTGTTGTTGCTGAATTCTTGAGCAATACAATTTTCAAGTATTGGTT<br>AAAGGGTCCGAAGGCAAACACTTATGAAGCCTTACTGACAATCGGGGGCCCA<br>GGGAACATCAAGAGGACTAAAGCTGGAGATTATTGGGGAGCCTCAAGTGATAA<br>TGTTGGAACCAAAGTTATCACAAGCGGTATAAGGTTTGATGAATCTGGCAAAAT<br>TTTAGAAGTCGTGCGTATCCCTCCACCGTATCAAGGCGAACATTTCGAACAAG<br>TTCATGAGCATAATGGGGCACTTTATATCGGGACTTTGTTTCACGACTTTGTAG<br>GTATATTGCACAATTACCAGGGTGCATCTGATCTGAAGGGAAATAATCTAGCTG<br>GGGCCAATGGAGCTTTGAATGGAGTGTCTGCTTCTGCCTGA |

| | |
|---|---|
| *OpSTR*<br>AB060341.1 | ATGCATAGTTCAGAAGCCATGGTTGTGTCGATTCTTTGTGCCCTTTTCTTGTCC<br>TCTCTTTCCCTTGTTTCTTCGTCGCCGGAGTTCTTCGAATTTATTGAAGCACCA<br>TCCTACGGACCAAACGCATACGCTTTTGACTCCGATGGTGAACTTTACGCTTC<br>CGTCGAAGATGGTAGAATTATCAAGTATGATAAGCCAAGCAACAAGTTCTTGAC<br>CCACGCTGTTGCCTCTCCAATCTGGAATAATGCACTTTGTGAGAACAATACTAA<br>CCAAGATCTAAAACCCTTGTGTGGGAGAGTATATGACTTTGGATTCCACTATGA<br>AACTCAGAGATTATACATAGCCGATTGTTATTTTGGTCTTGGTTTTGTTGGACC<br>CGACGGAGGCCATGCAATTCAGCTTGCCACTAGTGGAGATGGAGTGGAGTTT<br>AAGTGGCTTTATGCTTTGGCTATAGACCAACAAGCTGGCTTTGTTTATGTCACA<br>GATGTGAGCACAAAATATGATGACAGAGGTGTTCAAGATATCATAAGGATAAAC<br>GACACAACAGGAAGATTAATCAAATATGATCCATCAACTGAGGAAGTTACAGTT<br>TTGATGAAAGGGCTAAATATACCAGGAGGTACAGAAGTCAGCAAAGATGGCTC<br>CTTTGTTCTGGTTGGCGAATTCGCCAGCCATAGAATTCTCAAGTATTGGCTTAA<br>GGGACCTAAAGCAAATACATCAGAATTCTTATTGAAAGTCAGAGGTCCAGGAA<br>ACATAAAGAGGACAAAGGATGGAGATTTTTGGGTGGCCTCAAGTGACAATAAT<br>GGGATCACTGTTACTCCAAGAGGGATAAGGTTTGATGAATTTGGCAACATTTTA<br>GAAGTTGTGGCTATTCCTCTACCGTATAAAGGTGAACATATTGAACAAGTTCAA<br>GAGCATGATGGTGCATTATTTGTTGGATCTTTGTTCCACGAATTTGTTGGGATA<br>CTACACAATTACAAGAGTTCTGTTGATCATCATCAGGAGAAAAATTCAGGTGGG<br>CTCAATGCATCATTCAAGGAATTTTCTTCTTTCTGA |
| *OpSTR_2M* | ATGCATAGTTCAGAAGCCATGGTTGTGTCGATTCTTTGTGCCCTTTTCTTGTCC<br>TCTCTTTCCCTTGTTTCTTCGTCGCCGGAGTTCTTCGAATTTATTGAAGCACCA<br>TCCTACGGACCAAACGCATACGCTTTTGACTCCGATGGTGAACTTTACGCTTC<br>CGTCGAAGATGGTAGAATTATCAAGTATGATAAGCCAAGCAACAAGTTCTTGAC<br>CCACGCTGTTGCCTCTCCAATCTGGAATAATGCACTTTGTGAGAACAATACTAA<br>CCAAGATCTAAAACCCTTGTGTGGGAGAGTATATGACTTTGGATTCCACTATGA<br>AACTCAGAGATTATACATAGCCGATTGTTATTTTGGTCTTGGTTTTGTTGGACC<br>CGACGGAGGCCATGCAATTCAGCTTGCCACTAGTGGAGATGGAGTGGAGTTT<br>AAGTGGCTTTATGCTTTGGCTATAGACCAACAAGCTGGCTTTGTTTATGTCACA<br>GATGTGAGCACAAAATATGATGACAGAGGTGTTCAAGATATCATAAGGATAAAC<br>GACACAACAGGAAGATTAATCAAATATGATCCATCAACTGAGGAAGTTACAGTT<br>TTGATGAAAGGGCTAAATATACCAGGAGGTACAGAAGTCAGCAAAGATGGCTC<br>CTTTGTTCTGGTTGGCGAATTCGCCAGCCATAGAATTCTCAAGTATTGGCTTAA<br>GGGACCTAAAGCAAATACATCAGAATTCTTATTGAAAGTCAGAGGTCCAGGAA<br>ACATAAAGAGGACAAAGGATGGAGATTTTTGGGTGGCCTCAAGTGACAATAAT<br>GGGATCACTGTTACTCCAAGAGGGATAAGGTTTGATGAATTTGGCAACATTTTA<br>GAAGTTGTGGCTATTCCTCTACCGTATAAAGGTGAATTTATTGAACAAGTTCAA<br>GAGCATGATGGTGCATTATTTGTTGGATCTTTGCACCACGAATTTGTTGGGATA<br>CTACACAATTACAAGAGTTCTGTTGATCATCATCAGGAGAAAAATTCAGGTGGG<br>CTCAATGCATCATTCAAGGAATTTTCTTCTTTCTGA |
| *OpSTR_3M* | ATGCATAGTTCAGAAGCCATGGTTGTGTCGATTCTTTGTGCCCTTTTCTTGTCC<br>TCTCTTTCCCTTGTTTCTTCGTCGCCGGAGTTCTTCGAATTTATTGAAGCACCA<br>TCCTACGGACCAAACGCATACGCTTTTGACTCCGATGGTGAACTTTACGCTTC<br>CGTCGAAGATGGTAGAATTATCAAGTATGATAAGCCAAGCAACAAGTTCTTGAC<br>CCACGCTGTTGCCTCTCCAATCTGGAATAATGCACTTTGTGAGAACAATACTAA<br>CCAAGATCTAAAACCCTTGTGTGGGAGAGTATATGACTTTGGATTCCACTATGA<br>AACTCAGAGATTATACATAGCCGATTGTTATTTTGGTCTTGGTTTTGTTGGACC<br>CGACGGAGGCCATGCAATTCAGCTTGCCACTAGTGGAGATGGAGTGGAGTTT<br>AAGTGGCTTTATGCTTTGGCTATAGACCAACAAGCTGGCTTTGTTTATGTCACA<br>GATGTGAGCACAAAATATGATGACAGAGGTGGTCAAGATATCATAAGGATAAA<br>CGACACAACAGGAAGATTAATCAAATATGATCCATCAACTGAGGAAGTTACAGT<br>TTTGATGAAAGGGCTAAATATACCAGGAGGTACAGAAGTCAGCAAAGATGGCT<br>CCTTTGTTCTGGTTGGCGAATTCGCCAGCCATAGAATTCTCAAGTATTGGCTTA<br>AGGGACCTAAAGCAAATACATCAGAATTCTTATTGAAAGTCAGAGGTCCAGGA<br>AACATAAAGAGGACAAAGGATGGAGATTTTTGGGTGGCCTCAAGTGACAATAA<br>TGGGATCACTGTTACTCCAAGAGGGATAAGGTTTGATGAATTTGGCAACATTTT<br>AGAAGTTGTGGCTATTCCTCTACCGTATAAAGGTGAATTTATTGAACAAGTTCA<br>AGAGCATGATGGTGCATTATTTGTTGGATCTTTGCACCACGAATTTGTTGGGAT<br>ACTACACAATTACAAGAGTTCTGTTGATCATCATCAGGAGAAAAATTCAGGTGG<br>GCTCAATGCATCATTCAAGGAATTTTCTTCTTTCTGA |
| *OpSTR_8M* | ATGCATAGTTCAGAAGCCATGGTTGTGTCGATTCTTTGTGCCCTTTTCTTGTCC<br>TCTCTTTCCCTTGTTTCTTCGTCGCCGGAGTTCTTCGAATTTATTGAAGCACCA<br>TCCTACGGACCAAACGCATACGCTTTTGACTCCGATGGTGAACTTTACGCTTC |

| | |
|---|---|
| | CGTCGAAGATGGTAGAATTATCAAGTATGATAAGCCAAGCAACAAGTTCTTGAC<br>CCACGCTGTTGCCTCTCCAATCTGGAATAATGCACTTTGTGAGAACAATACTAA<br>CCAAGATCTAAAACCCTTGTGTGGGAGAGTATATGACTTTGGATTCCACTATGA<br>AACTCAGAGATTATACATAGCCGATTGTTATTTTGGTCTTGGTTTTGTTGGACC<br>CGACGGAGGCCATGCAATTCAGCTTGCCACTAGTGGAGATGGAGTGGAGTTT<br>AAGTGGCTTTATGCTTTGGCTATAGACCAACAAGCTGGCTTTGTTTATGTCACA<br>GATGTGAGCACAAAATATGATGACAGAGGTGGTCAAGATATGGTAAGGATAAA<br>CGACACAACAGGAAGATTAATCAAATATGATCCATCAACTGAGGAAGTTACAGT<br>TTTGATGAAAGGGCTAAATATAGCAGCAGGTACAGAAGTCAGCAAAGATGGCT<br>CCTTTGTTCTGGTTGGCGAATACGCCAGCCATAGAATTCTCAAGTATTGGCTTA<br>AGGGACCTAAAGCAAATACATCAGAATTCTTATTGAAAGTCAGAGGTCCAGGA<br>AACATAAAGAGGACAAAGGATGGAGATTTTTGGGTGGCCTCAAGTGACAATAA<br>TGGGATCACTGTTACTCCAAGAGGGATAAGGTTTGATGAATTTGGCAACATTTT<br>AGAAGTTGTGGCTATTCCTCTACCGTATAAAGGTGAATTTATTGAACAAGTTCA<br>AGAGCATGATGGTGCATTATTTGTTGGATCTTTGCACCACGAATTTGTTGGGAT<br>ACTACACAATTACAAGAGTTCTGTTGATCATCATCAGGAGAAAAATTCAGGTGG<br>GCTCAATGCATCATTCAAGGAATTTTCTTCTTTCTGA |
| *RsSTR*<br>Y00756.1 | ATGGCCAAACTTTCTGATTCGCAAACTATGGCACTGTTCACCGTCTTCCTTCTT<br>TTCCTCTCCTCTTCGCTCGCTCTCTCCTCTCCAATCTTGAAAGAGATTTTGATT<br>GAGGCTCCTTCCTATGCCCCCAATTCCTTCACCTTCGACTCAACCAACAAAGG<br>GTTCTACACCTCCGTCCAAGATGGCCGAGTTATCAAGTACGAAGGACCCAACT<br>CCGGTTTCGTCGACTTCGCCTATGCATCTCCCTACTGGAACAAAGCGTTCTGT<br>GAGAACAGCACAGATGCAGAGAAAAGACCCTTGTGTGGGAGGACATATGATAT<br>TTCATATAACTTGCAAAACAACCAGCTTTACATTGTTGATTGCTATTATCATCTT<br>TCTGTGGTTGGTTCTGAAGGTGGGCATGCTACCCAACTCGCCACCAGCGTTGA<br>TGGAGTGCCATTCAAGTGGCTCTATGCAGTAACAGTTGATCAGAGAACTGGGA<br>TTGTTTACTTCACCGATGTTAGCACCTTATATGATGACAGAGGTGTCCAACAAA<br>TTATGGATACAAGCGATAAAACAGGAAGACTAATAAAGTATGATCCCTCCACCA<br>AAGAAACAACACTACTGTTGAAAGAGCTACACGTTCCAGGTGGCGCAGAAGTC<br>AGTGCAGATAGCTCCTTTGTTCTTGTGGCTGAGTTTTTGAGCCATCAAATTGTC<br>AAATATTGGCTAGAAGGGCCTAAGAAGGGCACTGCGGAGGTTTTAGTGAAAAT<br>CCCAAACCCAGGAAATATAAAGAGGAACGCTGATGGACATTTTTGGGTTTCCT<br>CAAGTGAAGAATTAGATGGAAATATGCACGGAAGAGTTGATCCTAAAGGAATA<br>AAATTTGATGAGTTTGGGAACATTCTTGAAGTTATCCCACTCCCACCACCATTT<br>GCAGGTGAACACTTCGAACAAATTCAAGAGCATGATGGTTTGCTGTACATTGG<br>AACCCTGTTCCATGGCTCTGTGGGCATATTAGTATATGATAAGAAGGGAAATTC<br>TTTTGTTTCAAGTCATTAA |
| *RsSTR_2M* | ATGGCCAAACTTTCTGATTCGCAAACTATGGCACTGTTCACCGTCTTCCTTCTT<br>TTCCTCTCCTCTTCGCTCGCTCTCTCCTCTCCAATCTTGAAAGAGATTTTGATT<br>GAGGCTCCTTCCTATGCCCCCAATTCCTTCACCTTCGACTCAACCAACAAAGG<br>GTTCTACACCTCCGTCCAAGATGGCCGAGTTATCAAGTACGAAGGACCCAACT<br>CCGGTTTCGTCGACTTCGCCTATGCATCTCCCTACTGGAACAAAGCGTTCTGT<br>GAGAACAGCACAGATGCAGAGAAAAGACCCTTGTGTGGGAGGACATATGATAT<br>TTCATATAACTTGCAAAACAACCAGCTTTACATTGTTGATTGCTATTATCATCTT<br>TCTGTGGTTGGTTCTGAAGGTGGGCATGCTACCCAACTCGCCACCAGCGTTGA<br>TGGAGTGCCATTCAAGTGGCTCTATGCAGTAACAGTTGATCAGAGAACTGGGA<br>TTGTTTACTTCACCGATGTTAGCACCTTATATGATGACAGAGGTGTCCAACAAA<br>TTATGGATACAAGCGATAAAACAGGAAGACTAATAAAGTATGATCCCTCCACCA<br>AAGAAACAACACTACTGTTGAAAGAGCTACACGTTCCAGGTGGCGCAGAAGTC<br>AGTGCAGATAGCTCCTTTGTTCTTGTGGCTGAGTTTTTGAGCCATCAAATTGTC<br>AAATATTGGCTAGAAGGGCCTAAGAAGGGCACTGCGGAGGTTTTAGTGAAAAT<br>CCCAAACCCAGGAAATATAAAGAGGAACGCTGATGGACATTTTTGGGTTTCCT<br>CAAGTGAAGAATTAGATGGAAATATGCACGGAAGAGTTGATCCTAAAGGAATA<br>AAATTTGATGAGTTTGGGAACATTCTTGAAGTTATCCCACTCCCACCACCATTT<br>GCAGGTGAATTTTTCGAACAAATTCAAGAGCATGATGGTTTGCTGTACATTGGA<br>ACCCTGCACCATGGCTCTGTGGGCATATTAGTATATGATAAGAAGGGAAATTC<br>TTTTGTTTCAAGTCATTAA |

**Table S3. Sequences used to construct the phylogenetic tree.**

| Species and accession number | Sequence |
|---|---|
| *Camptotheca acuminata*<br>AES93117.1 | MAILKSSRTSAMLSIFISYFSCSSVLLVTASFQALPSPAPGPASFTFDLPLGI<br>GALYTGLADGRIVRYQRLRSTFVNYGYTAPNRNQAFCDGTNNTFLAPICGR<br>PLGLAFQFGTRRLYAADAAFGLVVIEPYGGPATQLATGVDGVRFRYPAAVD<br>VDQFSGTVYFTDASTRFNLSQLSQLIRTRDTTGRLLKYDPNTRQVTVLLRG<br>LAGPFAVAISSDRTYVLISEFIRNRIQKYWLTGPNANTAEVLLNVAGSPGNIR<br>RTIRGDFWVAINVQTPTVVLRGQRINGDGTILQTETFSPDFNTTLITEVNEY<br>GGALYLGSLYPKFVGVYKP |
| *Catharanthus roseus*<br>P18417.2 | MANFSESKSMMAVFFMFFLLLLSSSSSSSSSSPILKKIFIESPSYAPNAFTFD<br>STDKGFYTSVQDGRVIKYEGPNSGFTDFAYASPFWNKAFCENSTDPEKRP<br>LCGRTYDISYDYKNSQMYIVDGHYHLCVVGKEGGYATQLATSVQGVPFKW<br>LYAVTVDQRTGIVYFTDVSSIHDDSPEGVEEIMNTSDRTGRLMKYDPSTKE<br>TTLLLKELHVPGGAEISADGSFVVVAEFLSNRIVKYWLEGPKKGSAEFLVTIP<br>NPGNIKRNSDGHFWVSSSEELDGGQHGRVVSRGIKFDGFGNILQVIPLPPP<br>YEGEHFEQIQEHDGLLYIGSLFHSSVGILVYDDHDNKGNSYVSS |
| *Cinchona pubescens*<br>PQ568387.1 | MHISENMFVVTISFILFLSSPSLVLSSPYFQFIQAPSYGPNAYAFDSAGGLYA<br>VVEDGRIVKYEGSSNAFLDHAVASPFWTKKLCENNTKPQLKPLCGRAYDL<br>GFHYETQQLYIADCYFGLGVVGPEGGLAKKLAKSGDGVEFKWLYALVVDQ<br>QTGFVYVTDVSTKYDDRGVQDILRTNDTTGRLIKYDPTTREVTVLMKGLNV<br>PGGAEISKDGSFILIGEFLSNQILKYWLKGPKANTLEFLLHVKGPGSIRRTKA<br>GDFWVASSDNNGITVTPRGIRFDESGNILEVVPIPLPYKGEHIEQVQEHNGA<br>LYIGSLFHGFIGILYNYKGLSEENNLGGVVESLKGESFSF |
| *Coffea arabica*<br>XP 027088110.1 | MVKTMTKLIHVILIFSLAYVVRSDRTLNTFKILHASGPEAIAFDLTGQGPYTG<br>VSDGRVLKYEGPGIGFVEFAHASPLRTKEKCDGTDDPNLGPICGRPFGVG<br>FNYRTGELYIADAFFGLCKVGPDGGLAEQLATSAEGGPFKWLDGLDVDST<br>TEMVYFTDISTKYTFRELPQALSSGDSTGRLMSYNPKTKEVQVLLSGLQIP<br>GGTGVSRDGSFVLVSEYTGHRILKYWLKGPKANTAEAILSIHRPDNIKRTLL<br>GDFWIGANLIMQQPAPNTIPQGVRINESGKVLETINLDGIFRNETIAEVQEFA<br>GSYYVVSIRRNSVGICKCFVEENGRKLLGIRYDWIKENCS |
| *Cornus florida*<br>XP 059639506.1 | MAVILSFKAVAMLSIFILVICLPFSMGLFYPAFNKLQLPSTSIGPVSSAFDLFG<br>GGPYVGISDGRIVKYQGPPTNTFIDYAFDTPNRSKAVCDGTSNPDMGPLC<br>GRPLGLGFNYRTGELYITDAYYGLMVAGTNGRLATQLATSAEGVPFRFLTG<br>LDIDPTGIVYFADANTVYSLREIQQAIQSGDATGRLMKYDPNTKQVTVLLRG<br>LAFATGVAVGRNGGFVLVSEFKGNRIQKYWLKGPKANTAEILLNLPGPGIIK<br>RTTSGDFWVAANVLQQQPAQTAVRTGQRINEFGNVVQNVTLDAQYGNDLI<br>TEVQERLGTLYIGSLIKNFVGVLSPV |
| *Gelsemium sempervirens*<br>AXK92563.1 | MESLCVSQTMAVLSILVLFLISPPLVLSSCFSFLGPPSYGPNAYAWDWADE<br>GPYVAVEDGRIVKYEGPDINFVDFAYASPFWNKEQCLNNTSPDKKDLCGR<br>TYDLAFNYVTKELYVADCYFHLSVVGPEGGHATQLAKSVNGVPFKWLYAL<br>TVDQPTGNLYFTDVSTKYDDKGIQDIIRTKDTTGRVLKYDPSTKEVTLLMKD<br>LHVPGGIEVSKDSSFILVAEFMTHRILKYWLTGPKADTAEVLLKVRGPGNIK<br>RKITGEFWVASSDNNGITVTPRGILFDEFGRVLKVVPIPLPYKGEHIEQVVE<br>HDGKLYVGSLFHSYVGILQNFNHDDVKDKKSYSY |
| *Herrania umbratica*<br>XP 021287075.1 | MAYIISLRKVTLLSIFIFIFCSPSMVLSQFFSSIQLPPTARGPESFAFELGTGR<br>FYVGVDDGRILQYNGPATGFVDFGYTSGTRSKAVCDGATNPDLGPVCGRP<br>LGLGFHYASSQLYVCDAYVGLVSLDSGGGLANLVSSSAEGEPYRFCNGLD<br>VHQLSGNVFFTDTSTVYDLRNASKGLTANDSTGRLLKYDPNTKRVTVLLKN<br>LTGPAGAAVSQDGTYVLVSNFNSNTTMRYWLQGLRANTYDIISTQARPNNI<br>QGTLAGDFWQAAAMVKQPTQSLVPIGQRINGFGIVTRTVNFEPWYGNNLIS<br>GVQEFGGALYVASRYVNFVGVYSF |
| *Mangifera indica*<br>XP 044466508.1 | MDASVDIETDTCMQRSPNSTEIETLHICKHALQNTCSHFVHFHYFLLVFAFC<br>SSFFPISKAATPFSRIRPESSAFDAAGGGPYTGVADGRILKYLNQVDGFVEY<br>GVTSRNWSRALCVSNINPALGPTCGRPIGLGFNNFTGSLNIADAYLGLLELP<br>ANQTVATQLATSADGVPFRFPDGLDVDPLTGDVYFTDASAVYQISQLQLAV<br>ALNDSTGRLLKYDIKTKRVTVLLKGLAGAAGTAVSSDGSFVLVTEFIGQRIQ<br>KYWLKGPKANTTEVIITFHGRPDNIKRNPGGDFWVAVNQKTLIPLIMLPEGI<br>RLDANGTIRESLPISVQYGLQSISEVQQFSGKLYIGSLYGNFIGTYE |

| | |
|---|---|
| *Mercurialis annua*<br>XP 050238081.1 | MQYQSDRLRKICEVSLIFMASMSLIFILLLHVHFAMAHPRASFERLYLPTPLV<br>GPESLAFDWNGGGPYTGVSDGRILKYQGPNLNFTEFAYTTQNRNKSQCD<br>GATDPSLQSICGRPLGIAFYYRTGELYIADANNGLFVVGPNGGQATRLLNS<br>VQGVPLKFLAGLDVDQTTGNVFFTEASSAFQLKDIVGLVQSRDNTGSLFMY<br>DPRTKRATVLLRNLAAAAGTAVSRDGSFVLVSEFLANRIQKYWLRGPRANT<br>AQILRSFRGKPDNIKRNANGQFWVAVSVLRDPPPPPKPRMLPLGYRINENG<br>MVLQVVSFRGPYETEAISEVQEMNGTLYAGSLHVNFAATMFRR |
| *Mitragyna speciosa*<br>ABZ79473.1 | MNTSESMVALTIFFALFLSPLSVVLSSAEFFQFLKSPYGPNAFAFNSAGELY<br>AAVEDGRIVKYKGSSNHGFSTHAVASPFWNRKVCENYTELQLKPFCGRTY<br>DLGFHYETQQLYIADCYYGLGVVGPEGGRATQVARSADGVDFKWLYALAV<br>DQQTGFVYLTDVSIKYDDRGVQDILRINDTTGRLIKYDPSTNEARVLMNGLN<br>VPGGTEVSKDGSFLVVAEFLSHRILKYWLKGPKANTSEVLLKVRGPGNIKR<br>TKAGEFWVASSDNNGITVTPRAIKFDDFGNILQVVPVPPPYKGEHFEQAQE<br>HNGSLYIGTLFHDFVGILHNYEGSSDPKENNVDGVDGSLNGVASSV |
| *Mitragyna speciosa*<br>TRINITY_DN16400_c0_g<br>2_i1 | MDILFILILLVSSTSLVLSSPLLKSIQSPYGPNAFAFDTEGQLYSAVEDGRIVK<br>YERSFTGFVDHAYASPFWNKQLCQCKNTTLEAHLKPLCGRAYDLGFNHAT<br>NQLFIADGYYGLGVVGPEGGQAVQLASSAEGVDFKWLYALTVDQRTGIIYF<br>TDVSTIYDDRGVQDIIDTNDTTGRLLKYDPSTKEVTVLLKGLNVPGGTEVSN<br>DSSFVLVGEYLSNKILKYWLKGPKANSSEILLKIPGPGNIKRTARGDFWVPS<br>TDDNAPILTSIGVRFNEFGRILETVDLPQPYKTEHLEQIHEHNHALCVGSLFH<br>NFIGVLERASLDQQKEHGLNGLNEYTKESASD |
| *Nothapodytes nimmoniana*<br>ASY08091.1 | MSSKLLLASVTIFIISCLLSSVLKFSENLYKSSSSSSSSSNEEEEVNNIPIDEG<br>AFGPESFAFDPSGGGPYTGVSDGRIIKWQPHQRRWINFATTSSQRHGCER<br>TSAQQVSEHICGRPLGLVFNQKSGDLYIADAYMGLLSVGPNGGLANQIATQ<br>AQGIPLAFTNALDIDQSNGLVYFTDSSSVYKRRNYVSAIVSGDKTGRLMKY<br>DPKTSEVTVILKNLSFPNGVSLSRDGDYILIAETTNCRILKFWIKTSKAGMVE<br>VLAELPGFPDNIKRNEKGEFWVGIYGRRGNFIKWVLSSPWIGNILVKLPLDIT<br>EIYSYLASFRACGLAVKLNGEGKILVLLEDENGKKWKFISEV |
| *Ophiorrhiza pumila*<br>Opuchr05 g0008180-1.1 | MAIILTLILFLSSTVDVLSSPILNQITSPYGPNAFAFDSAGRLYSAVEDGRIIKF<br>ERSGNTFVDHAVASPLWNKTVCKAKHTYVESHLKPLCGRVYDLGFSYAEN<br>QLYIADGYYGLGVVGPDGGCAIQLANTADGVDFKWLYALTVDQLTGIVYFT<br>DVSTIYDDRGVQDILDTNDTTGRLLKYDPSTKEATVLLKGLNVPGGTEVSK<br>DSSFVLVGEYLSDRIRKYWLKGPKANTSEILLEIPGPGNIKRTKNGDFWVPS<br>TDDKSPIITSVGVRFDGFGRILETVDLPQPYANEHLEQINEHNGALFIGSLFH<br>SFIGVLQKDFLAAHDKEKPSSPY |
| *Ophiorrhiza pumila*<br>Q94LW9 | MHSSEAMVVSILCALFLSSLSLVSSSPEFFEFIEAPSYGPNAYAFDSDGELY<br>ASVEDGRIIKYDKPSNKFLTHAVASPIWNNALCENNTNQDLKPLCGRVYDF<br>GFHYETQRLYIADCYFGLGFVGPDGGHAIQLATSGDGVEFKWLYALAIDQQ<br>AGFVYVTDVSTKYDDRGVQDIIRINDTTGRLIKYDPSTEEVTVLMKGLNIPG<br>GTEVSKDGSFVLVGEFASHRILKYWLKGPKANTSEFLLKVRGPGNIKRTKD<br>GDFWVASSDNNGITVTPRGIRFDEFGNILEVVAIPLPYKGEHIEQVQEHDGA<br>LFVGSLFHEFVGILHNYKSSVDHHQEKNSGGLNASFKEFSSF |
| *Pistacia vera*<br>XP 031276814.1 | MDSIFAPAAMFPISLIIFFLSLPSVLHSLNYVKLPLPLFAVGPESSAFDAAGAG<br>PYTGVADGRILKYVNPSVGFVDYGVTSPNRSKALCVGYSIPALGPTCGRPL<br>GLGFNNLTGDLYIADAYLGLLVLPKNQTIATQLATSANGVPFRSPDGLDVDP<br>VTGDVYFTDASAVYRISQILLAIAKNDSTGRLLKYDIKTKQVTVLLKGLSGPA<br>GTAVSPDGTFVLVTEFIGKRIQKYWLKGPKANTAEVLITFKGRPDNIKRNRG<br>GDFWVAVNQETIIPLVMKPEGIRLDANGTVKESRPINLQYGTTLVSEVQEFS<br>GALYIGSMYTNFIGVYK |
| *Pogonopus speciosus*<br>Epi-STR | MKSQSLANMRSSESMLAVSIFLVLFSSSPSLVLSSPLFFQFLKSPFAPNAYG<br>FGASGELYASVEDGRIVKYEGSSNRFLTHAVASPHWKRDLCENNTNPQLK<br>PFCGRIYDFGFNYGTQQLYIADSYYGLCVVGAKGGRATQLVTSADGVDFK<br>WLYGLALDQHTGIVYFTDVSSVYDDRGGQDMVRTKDKTGRLLKYDPSTDE<br>VTVLMKELGIAAGTEVSEDGSFVVVAEYLSNTIFKYWLKGPKANTYEALLTI<br>GGPGNIKRTKAGDYWGASSDNVGTKVITSGIRFDESGKILEVVRIPPPYQG<br>EFFEQVHEHNGALYIGTLHHDFVGILHNYQGASDLKGNNLAGANGALNGV<br>SASA |
| *Prosopis alba*<br>XP 028753725.1 | MSNIAILVVFALLICSSSVANSNRLLSRQFLPPPLTGPESLAFDSVGEGPYTG<br>ASDGRILKYVGPSDGFVEFASSSPCRNKTICDGISDFSEIQKTCGRPLGLAF<br>NYQTSELYIADAYFGLMKVPSSGGSPTQVVSCVEGKSFRFLAGLDIDPQTG<br>VVYFTEASTTYQIRDLPTLLRSGDNSGSLVKFDPCTMETSVLLRDLAVPSG<br>VAVSKDGSYVLVSEFMANRVQRFWLEGDKQNTSEIFLQLPGKPDNIKRNS |

| | |
|---|---|
| | AGDFWVAVNNQVGSPPPSRPPVLPLAVRVDGNGFVLQEVALVEEYGTEM<br>VSEVQEFNGKLYSASLDVSYVNVFMP |
| *Rauvolfia serpentina*<br>P68175.1 | MAKLSDSQTMALFTVFLLFLSSSLALSSPILKEILIEAPSYAPNSFTFDSTNKG<br>FYTSVQDGRVIKYEGPNSGFVDFAYASPYWNKAFCENSTDAEKRPLCGRT<br>YDISYNLQNNQLYIVDCYYHLSVVGSEGGHATQLATSVDGVPFKWLYAVTV<br>DQRTGIVYFTDVSTLYDDRGVQQIMDTSDKTGRLIKYDPSTKETTLLLKELH<br>VPGGAEVSADSSFVLVAEFLSHQIVKYWLEGPKKGTAEVLVKIPNPGNIKR<br>NADGHFWVSSSEELDGNMHGRVDPKGIKFDEFGNILEVIPLPPPFAGEHFE<br>QIQEHDGLLYIGTLFHGSVGILVYDKKGNSFVSSH |
| *Rehmannia glutinosa*<br>KAK6133690.1 | MVPFLLLLFCLPNRALANGFRSFKMIPLPSHGCEAYAFDSDNGGPYTGLND<br>GRIVKYQGPKIGFVEFATTVPNRSKELCDGKNGDDPKTGPLCGRPIGLEFN<br>HRTGELYVADAFRGLMVVARGGGVAARLAGGRDGVPFDAPDAIAIDPITGE<br>VYFTDVGSIFFKTTNMTEILLSGDTSGRLLKYDPKTKQRTVVLTGLAVPNGV<br>AVSKDGSFVLVAEYIACRITRFWLKGPQANTSDIFVQLPGNPDNIKRTKSGD<br>FWVPVNIQKLYPKLISFPLGQKINARGEIIETVNFYAEYNATYITEVHEQLGSL<br>YVASVYTNFVGVYRGLKCSSSGIAHPL |
| *Rosa chinensis*<br>XP 024196605.1 | MKLYFLSLLFFFSLILASSTPASAFFVVNLTNYHQIELPKSVVGPESIAFDCR<br>GKGPYVGVSDGRILKWQGPCRGWTEFAFTSPTRPRKICDGSTDNTTEPIC<br>GRPLGLKFNPKTCELYIADAYFGLLKIGRNGGRPQQLATSLNGVPFRFLNAL<br>DIDSQSGIVYFTDTSTVYQRRLWPLSIATGDKTGRLLQYDPCTKKVTVLLDN<br>LAFANGVALSEDSSFLLVAESATFKIFKLWLRGPKAYDVKLFTQLKRPADNI<br>KRTNGGEFWAALNSLRGLQGNQTSPSLWLTKDPVGVKFDEQGNVMEVLD<br>GEGGPTLESISEVEEHNGKLWIGSVVKSYVGVIKS |
| *Strychnos nux-vomica*<br>Cluster-17159.0 | MRSSTHTMAVGLTISILLLASSSLALSSLFLELIDSPYGPNAFTFDSQDRLYA<br>AVEDGRIVRYEGPSAGFVDFAYASPFWNKSLCGNNRNPAKKPLCGRTYDL<br>AFHHATKQLYLADCYYTLSVVGSEGGHAKRLVKSANGVPFKWLYALTVDQ<br>NTGFIYFTDVSTIYDDRGVQEIINTRDHTGRLLQYDPLTDEVTVLMNGLYVP<br>GGTEVSQDSSFVVVGEFLTHRILKYWLKGPKANTAELLLRVRGPGNIKRTT<br>SGDFWVASSDNNGITVTPRAIRFDDFGNILQVVPVPPPYKGQHIEQVQENN<br>GILYIGSLFHSFVGKLYGISDEEKRENIDGVNIYFRGSSSS |
| *Tabernaemontana elegans*<br>AEY82398.1 | MANSQIVAVFTIFLLFLSSPSLALSSSILGEIQIDAPSYGPNAFTFDSTNKGFY<br>TSVLDGRVLKYDGPETGFVDFAYASPYWNKAFCENNTDAEKRPFCGRAY<br>DIAYGYKSNNLYIVDCYFHLSVVGPEGGHATQLSTSVEGVPFKWLYALAVD<br>QRTGLIYFTDVSTRYDDRGVEEIIMTSDRTGRLIKYDPSTKETTLLLKELHVP<br>GGAEVGADSTFVLVAEFLNDQILKYWLEGPKKGTAEVLLKIPKPGSIKRNAK<br>GHFWVASSEEQGGMHGIVTPRAVRFDEFGNILEVFLMPPPYAGEHFEQVQ<br>EHDGLLYVGTLFHSAVGILIYNEKGNSGVKGQ |
| *Tabernanthe iboga*<br>Unigene2910_All | MVNSNTKSQIMAAFTIFLLFLSSPSVALSSSILGEIQIDAPSYAPNAFTFDSTN<br>KGFYTSVQDGRILKYEGPETGFIDFAYASPYWNKAFCENNTDANKRPLCG<br>RAYDIAYDYKGNNLYIVDCYFHLSVVGPEGGHATQLTTSVEGVPFKWLYAL<br>SVDQRTGIIYFTDVSTIFDDRGVEEIMMTSDRTGRLIKYDPSTKETTLLLKEL<br>HVPGGAEVGADSSFVLVAEFLSDQILKYWLEGPKKGTAEVILKIPKPGNIKR<br>NAKGHFWVSSSEEQGGMHGIVTARAVRFDEFGNILEAFLMPPPYAGEHFE<br>QVQEHDGLLYVGTLFHSTVGILIYNEKGNSGVSGQ |
| *Theobroma cacao*<br>XP 007037730.2 | MAYIISLREATLFSIFIFIFCSPSMVLSQFFSSIQLPPTAKGPESFAFELGTGRI<br>YVGVDDGRILQYNGPATGFVDFGYTSGTRSKAVCDGATNPDLGPICGRPM<br>GLGFHYATSQLYVCDAYVGLVALDSRGGLANLVSSSADGEPYRFCNGLDV<br>HQLSGNVFFTDTSTVYDLRNASKGLTSNDSTGRLLRYDPNTKRVTVLLKNL<br>TGPAGAAVSQDGTYVLISNFNSNTTMRYWLQGPRANTYDIISIQARPNNIQ<br>GTLAGDFWQAAAMVKQPTQSLVPIGQRISGLGIVTRTVNFEQWYGNNLISE<br>VQEFGGALYVASRYVNFIGVYRF |
| *Tripterygium wilfordii*<br>XP 038688094.1 | MISLFVILFTFPSIILCNVTPSFGRLPLSPNALGPEALAFKRLGRGPYTGVED<br>GRILQYLPSAGFRDFAFTSPNRSNAVCDGNTNQSLGPVCGRPLGLGFYFQ<br>TGDLYIADAYRGLLVVGQNGGLAIQLAISADGVPFRFTDGLDVDQQTGIVYF<br>TDASSVYPRSAADEALARGDATGRLLKYDPRTAQVTVLLRGLQLASGTAVS<br>IDGSFVLVTEFGARRVQKYWLRGPRANTAEILVTLPGNPANIKRTLLGDFW<br>VAVNINTTTIRPTGIKINGLGKILRTVPLDALYNNIRVSEVQEFLGKLYIGSTTE<br>TFVGVVNI |
| *Uncaria rhynchophylla*<br>CNA0013842 33474 | MSSGLKLSMSMLTVFFLFFFSSSSFVLSAPILKSIQAPYGPNAFAFDSAGGL<br>YTAVEDGRVVKYGGPGIGFLNHAYASPFWNETLCKSKNTTIEAHVKPICGR<br>VYDLGFNYATEKLYIVDGYYGLGMVGPHGGLAIQVANGADGADFKWLYAL<br>TVDQRTGIVYFTEVSSVYDDRGVQDIMDTNDTTGRLLKYDPLTKKVTVLMK |

| | |
|---|---|
| | GLNVPGGTEVSKDSTFVIVGEYLGNKILKYWLEGPKANSSEILLEIPGPGNIK RTPAGDFWVPSTDENAPILTSIGVRFNEFGRILETVDLPQPYKNEHLEQIHE HNHALYVGSLFHNFIGVLERASLDQQKEYGLDGLDEYNKESFPY |
| *Uncaria rhynchophylla* CNA0013842 88809 | MLSEVRLNMHSSMVALTIFLILFWSPLSVVLSSAEFLQFIKSPYGPNAFAFN SAGELFATVEDGRIVKYKGKGPSNGFSTHAVASPVWNRKVCENNSKPELK PFCGRAYDLGFHYETQQLYIVDCYYGLGVVGPEGGRVTQLAKSADGRDFK WLYALAVDQQTGSLYTTDVSAKYDDRGVQDILRINDSTGRLIKYDPSANEA SVLLKDLNIPGGVELSQDGSFLLVGEYLSHRILKYWLKGPKANTWEVFFKV RFPGNIKRNNAGEFWIASSDYNGITVTPRGFRVDESANVLEVVPIPPPYKG EFYEQVQEHNGALYVGSLFHDFVGILHNYEGSSDPKENNADGFNGSLNGL ASSV |
| *Uncaria rhynchophylla* USE06681.1 | MVLLTITFILFLSPLSVVQSSPEFFQFLKSPYGPNAFAFNSAGELYAAVEDG RIVKYKGSNNGFSTHAVASPIWNGEVCENNTKPQLKPFCGRAYDLGFHYE TQQLYIADCYYGLGVVGSEGGHATQLARSADGVDFKWLYALALDQQTGFV YLTDVSTKYDDRGVQDILRINDTTGRLIKYDPSTNEARVLMNGLNVPGGTE VSRDGSFLVVAEYLSHRILKYWLKGPKANTSEVLLKVRGPGNIKRTHDGEF WVASSDNNGITVTPRGIKFDEFGNILEVVPIPLPYKGEHFEQVQEHNGALYI GSLFHDFVGILHNYEGSSDAQKNYIDGVNGSLNGEASLV |
| *Uncaria sinensis* CNA0013752 64143 | MSMLTVFFLFFFSSSSFVLSAPILKSIQAPYGPNAFAFDSAGGLYTAVEDGR VVKYGGSGIGFLNHAYASPFWNETLCKTKNTTIEAHLKPICGRVYDLGFNY ATKKLYIVDGYYGLGMVGSHGGLAIQVANGADGADFKWLYALTVDQRTGIV YFTEVSSVYDDRGVQDIIDTNDTTGRLLKYDPLPKKVTVLMKGLNVPGGTE VSKDSTFVLVAEYLGNKILKYWLEGPKANSSEILLEIPGPGNIKRTPAGDFW VPSTDENAPILTSIGVRFNEFGRILETVDLPQPYKNEHLEQIHEHNHALYVGT LFHNFIGVLERASLDQQKEYGLDGLNEYNKESFPY |
| *Vinca minor* AEY82399.1 | MALFTALLLLLSSSLVLSSPILKVILIDTPSYAPNSFTFDSTNKGFYTAVQDGR ILKYQGPNSGFIDFAYASPYWNRGLCEKTRDEEKKPICGRTYDIAYYYKKKE IYIVDSYYHLSVVGAEGGYSTQLSTSVDGVPFKWLYAVTVDQTTGLVYFTD VSSIYHDSPEGIEAIMGTSDRTGRLMRYDPSTKETTLLMKELHVPGGAEISA DSSFIVVAEFLSNRIVKYWLQGPKKGTTEFLVKIPNPGNIKRNKDGHFWVSS SEEEGGQHGKVTARGIKFDEFGNILQVILLPPPYVGEHFEQIQERDGLLYIG TLFHGSVGILQYYPEKGIPSVSSQ |
| *Vitis vinifera* XP 010663511.1 | MFYPLFLFFFLFSHGNLSFGDNSSSLFRSFRMLHLPTPTIGPEALAFDCSGA GPYASVADGRVLKWQAESAGFVDFTVASPSRSKQLCDGSSDPAKEPTCG RPLGIGFNNKTGDLYIADAYYGLFVVGPDGGRATQLATEAEGVPFRFLNAV DVDQETGIVYFTDASARFQRREFQNAVLAGDMTGRLMKYDPRTKQVTVLL RGLGLAVGVAINKDGSFVLVSEFIATRIQRYWLRGPKANTSELFLKPTGTPD NIKRNARGEFWVAANIGAEMAAAAPLGLRVSEEGKILQVVAFDTGDITRTIS EVHEYNGALYVGSLALPFVGVFTF |
| *Ziziphus jujuba* XP 024930052.2 | MLSILIFFFCFPSVVFSISNCKKIDLPLTAVGPESSTFDSWGQGPYTAISDGR VVKYNGPGVGYVDYVITSPFRSKRECDGTNNPELWPKCGRPIGIAFYFLKN QLFIADAYRGLLVAGPNERLATQIATSAEGLPFKFLDGLDVDQLTGNVYFTD ASSVYTLRQFAQAVAANDASGRLLKYDPKTGQVTVLQRGLSGAAGVAVSL DSSYVLVTEYIANRIQKFWLRGPKANTSEILVTLEGRPDNIKRTPAGNFLVA VTIQNASTQTIVATAVTINGVGNIRQSVSLGPPYDNTSISEVLKFRHSYYIGS LMADFLGVCN |
